# ClassiCOL: LC-MS/MS analysis for ancient species Classification via Collagen peptide ambiguation

**DOI:** 10.1101/2024.10.01.616034

**Authors:** Ian Engels, Alexandra Burnett, Prudence Robert, Camille Pironneau, Grégory Abrams, Robbin Bouwmeester, Peter Van der Plaetsen, Kévin Di Modica, Marcel Otte, Lawrence Guy Straus, Valentin Fischer, Fabrice Bray, Bart Mesuere, Isabelle De Groote, Dieter Deforce, Simon Daled, Maarten Dhaenens

## Abstract

LC-MS/MS extends on the MALDI-TOF ZooMS approach by providing fragmentation spectra for each peptide. However, ancient bone samples generate sparse datasets containing only a few collagen proteins, rendering target-decoy strategies unusable and increasing uncertainty in peptide annotation. ClassiCOL embraces and even extends this ambiguity using a novel ‘isoBLAST’ approach. The exhaustive set of potential peptide candidates created in this way is then used to retain or reject different potential paths at each taxonomic branching point down to the taxonomic level attainable with the sample information, always allowing for potential mixtures in the process. As an end point, all considered ambiguity is graphically represented with a clear prioritization of the species in the sample. Using public as well as in-house data, we demonstrate the performance of this universal postprocessing approach on different instruments and explore the possibility of identifying genetic as well as sample mixtures. Diet reconstruction from 40,000 year old cave hyena coprolites illustrates the exciting potential of this approach.

**Teaser:** ClassiCOL is a postprocessing tool that allows for more accurate species classification from LC-MS/MS measurements of collagen.

## Introduction

Bone morphology-based species classification has been the state of the art for both paleontological and zooarcheological research for decades. However, when specimens are degraded, fragmented, and/or fractured to a point where this methodology can no longer be used, paleoproteomics has proven to be an excellent candidate to tackle this challenge (*1*). Compared to DNA, proteins tend to survive longer, particularly in biomineralized matrices like bone, enamel or eggshells (*2*–*7*). Together with the low sample amount required, straightforward and increasingly automatable sample preparation and relatively low cost, this makes palaeoproteomics a very appealing approach (*8*–*11*).

When analyzing ancient bone samples, often only collagen proteins remain detectable, especially COL1A1/COL1A2, which are the most abundant and stable proteins, estimated to constitute over 90% of protein in bone, *in-vivo* (*5, 12*). This abundance inspired the first successful ancient protein methodologies relying on MALDI-TOF MS, to create species-specific peptide mass fingerprints (PMF) (*13*–*17*). This method is generally referred to as Zooarcheology by Mass Spectrometry (ZooMS), and can be performed at very high sample throughputs. More recently, methods have been adopted that use liquid chromatography tandem mass spectrometry (LC-MS/MS) for species determination because this provides intrinsically more information-rich data. We refer to such approaches as ZooMS^2^. For example, SPIN relies on curated peptide-spectrum matches (PSMs) to accurately classify species, using a custom database containing the most commonly found proteins in archeological mammalian bones (*8*).

Essentially, an LC-MS/MS system extends on the MALDI-TOF approach by measuring not only the intensity and *m/z* of the precursor masses but also providing retention time (t_R_) and a fragmentation pattern for each peptide (*18, 19*). Briefly, the spectra that are generated from peptides are translated into a simplified, i.e. peak-picked, list of ion coordinates including at least the precursor mass and fragment ions with their respective intensities (*20*). Peptide sequences are then obtained by scoring algorithms that perform a search against a translated DNA sequence database. Generally, these report the best peptide hit for each spectrum in the data (*21*). To threshold true identifications from false ones, a probabilistic approach was developed by Matrix Science (Mascot) (*22*), and later also in MaxQuant (*23, 24*).

As general MS data became increasingly complex, a strategy to empirically assess the false discovery rate (FDR) at a given score threshold (rather than computing it probabilistically) was established through the target-decoy approach (*25, 26*), as an alternative to algorithms like PeptideProphet (*27*) that fit the discriminant score distributions of true and false hits based on fixed weights of the different metrics or scores. To do so, the sequences in the database are reversed or scrambled into nonsense sequences and added to the database. When searching against this database, the score can be cut off at a threshold below which a given portion of decoy sequences (and thus false discoveries) have been identified, In a later stage, these score distributions of ‘true’ and especially ‘known false’ annotations became a means to improve the weights that were given to specific features of the scoring algorithm. The first algorithm using this was Percolator, the first widely adopted, support vector-based machine learning strategy in proteomics, still being used to date (*28, 29*). The further increase in data richness obtained through MS sparked the latest class of search engines with the introduction of more elaborate machine learning approaches, which rely on neural networks and accurate predictions to extract ever-larger numbers of peptide identifications from the data (*30, 31*). Still, the target-decoy approach is pivotal for all these algorithms, since they need to be trained on what is right and what is wrong in order to improve the feature weights used to score the peptide-to-spectrum matches (PSMs).

This presents paleoproteomics with a conundrum: ancient bone samples generate sparse datasets containing only a few hundred peptides derived from only a few proteins, effectively disabling the use and efficiency of target-decoy strategies for FDR estimation and potential score improvement. In other words, where the simplicity of the protein composition enables fast and effective PMF using MALDI, it actually complicates LC-MS/MS data interpretation. Fortunately, probabilistic search engines like Mascot and MaxQuant are still suitable for sparse peptide identification and are therefore the preferred choice for most proteomic challenges involving data sparsity. Still, the data consists of peptides that can be derived from the orthologous collagen sequences of a plethora of organisms, producing lists of very similar and often indistinguishable peptide candidates for each spectrum (*32, 33*). This ambiguity is not easily resolved because it is caused by positional isomers and isobaric changes that do not affect the score, such as posttranslational modification (PTM) combined with amino acid substitutions. This is further aggravated by the taphonomic processes that lead to low quality spectra (*34, 35*). In the end, this ambiguity impairs accurate post-processing tools like Unipept (*36*), which infers unique peptides to species at their last common ancestor (LCA), and leading to incorrect classifications by an otherwise highly performant tool.

As the end goal of zooarcheology by MS is to identify the species of origin rather than the proteins inside the sample, we propose here to embrace and even extend the ambiguity in MSMS search outputs to obtain an exhaustive set of potential peptide candidates (PPCs). This increases the chance of containing the correct peptide sequence for each spectrum in an otherwise redundant list of PPCs. This new axiom is then leveraged by rejecting the respective peptide sequences during species classification. Therefore, an anti-Occam’s razor (*37*) based algorithm is used to follow the NCBI taxonomic tree and, at every branching point, discard the branches that are mere subsets of the other and do not contribute unique peptides. We demonstrate the efficiency of this approach on several public datasets and in-house generated data, showing that this universal post-processing tool enables the identification of different species from single bone fragments as well as mixtures derived from their remains (e.g. from dust in bags or coprolites). Moreover, we show that this methodology holds equally high potential for identifying protein-containing paint and glue binders in cultural heritage samples.

## Results

### Establishing an extensively curated collagen database for ZooMS^2^

The most prominent obstacle in developing LC-MS/MS-based paleoproteomic approaches is the low number of peptides and their derived spectra, i.e. data sparseness, and in the case of ZooMS^2^, the exclusive focus on collagen proteins. We have compiled a database that is comprised of 221 species of Mammalia, supplemented by Reptilia (*n*=45), Osteichthyes (*n*=157), Chondrichthyes (*n*=11), Aves (*n*=162), Amphibia (*n*=15), and Cephalopoda (*n*=3), from which all collagen isoforms, up to 45 per species, were downloaded in FASTA format from both the UniProt and NCBI repositories **(Supplementary data 1)**. Hereby, the diversity of species searched is significantly extended compared to general ZooMS workflows, where this is still a limiting factor (*17*). This collagen database contains all available collagen proteins (beyond the conventional COL1A1 and COL1A2) and will be continuously maintained on the ClassiCOL Github page as new species are submitted to the repositories.

Importantly, this ClassiCOL DB is best deployed during the initial probabilistic database search with e.g. Mascot or MaxQuant. Still, CSV search result outputs from more constrained database searches can still be rescued through the isoBLAST ambiguation described below, albeit to a less precise taxonomic level.

### Mapping the ambiguity problem in conventional ZooMS^2^ searches

Collagen databases differ substantially from conventional proteome databases because of their highly similar tryptic peptide composition. When using databases consisting of highly similar protein sequences, conventional search engines like Mascot provide taxonomically ambiguous results similar to the more elaborately described protein inference problem (*38*) (**Figure 1**). Therefore, this equivalent “*species inference problem*” makes species classification very challenging.

**Figure 1.**
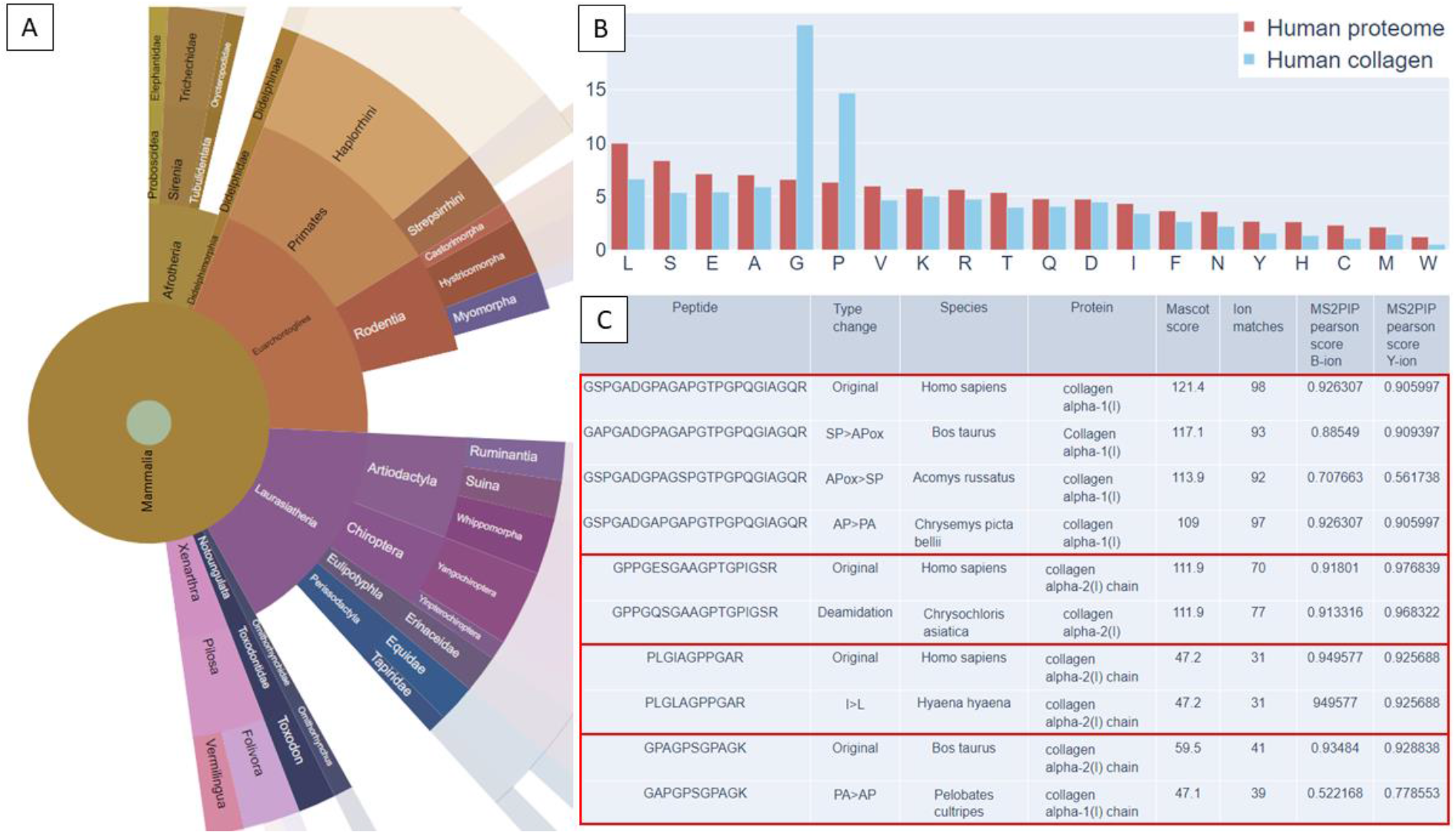
Demonstration of taxonomic uncertainties from a ZooMS^2^ output file. **A**. Unipept (*36*) sunburst view on Mascot result of *Homo sapiens* reference sample of Rüther et al, 2022 (*8*) showing uniqueness of peptide stretches on different taxonomic levels (**Supplementary figure 1**). Sunburst plot is colored via phylogenetic relatedness. **B**. Histogram showing the percentage amino acid distribution between the human reference proteome (UP000005640), collected from Uniprot (red) and solely collagen sequences extracted from the same human proteome from Uniprot (blue). **C**. Demonstrative overview of *Homo sapiens* collagen related peptides, alongside the top results with similar Mascot scoring. Due to an isobaric switch the protein of origin and species of origin can change. Unfortunately, intensity prediction and the accompanying MS^2^PIP (*40*) score does not always change either, compromising this second pass solution to further filter the peptide results.

Another layer of ambiguity is added when peptides cannot be distinguished from one another due to isobaric changes, which can lead to identification of false positive, species-specific, unique peptides (**Figure 1A**). In ZooMS^2^, isobaric changes include **(i)** a switch of two consecutive amino acids (positional isomer), **(ii)** a single residue isobaric switch including isoleucine-to-leucine (only resolvable in MS^n^ acquisitions), and N or Q deamidation versus D or E unmodified amino acids, **(iii)** mono-dipeptide changes e.g. AG to Q and **(iv)** combined PTM and amino acid switches, e.g. the isobaric alanine-to-serine substitution (+ 15.994915 Da) which can occur due to a nearby hydroxyproline (-PS-to -P_ox_A-). The chance of correctly identifying isobaric shifts further decreases when the general amino acid distribution in the database is greatly shifted compared to full proteomes, for which these search algorithms are designed. This is certainly the case for collagenous samples, which are strongly enriched in glycine and proline (**Figure 1B**).

This issue is profound, as many amino acid differences between species involve isobaric changes. This has been recognized by Buckley et al. (2019), who suggested that the isobaric shifts within peptide sequences could only be resolved by an in-depth investigation of the tandem spectra (*6*). Later, Rüther et al. (2022) (*8*) suggested to automate this process by calculating additional scores to filter the highest quality peptide matches, in contrast to Gilbert et al., (2024) (*39*) who have explored the capability of MS^3^, wherein MS/MS fragments are further fragmented. Alternatively, standard first-hit-export files have been used, yet these contain a random selection of correct and incorrect annotations making it futile to use well-established species inference post-processing algorithms such as Unipept (*36*). On the other hand, exports containing all ambiguous peptide-to-species matches completely impair the use of a species-specific peptide selection.

Generally, these isobaric changes will barely affect the (Mascot) scoring, especially if the resolving fragment is low in abundance and the scoring will not be affected if the resolving fragment is absent (**Figure 1C**). Still, recent advances in intensity prediction could theoretically help to prioritize the different options identified by the search engine. We verified this by comparing the Pearson correlation of the measured intensities of the fragments with an MS^2^PIP intensity prediction (*40*) for the original sequence and one of the aforementioned allowed isobaric changes by isoBLAST. **Figure 1C** shows how for several peptides from **Figure 1A** Mascot scores of isobaric peptides are very close and could easily switch ranks based on slight changes in the spectral quality. Unfortunately, even machine learning-based algorithms that accurately predict the y-ion and b-ion intensities cannot always resolve these isobaric changes in peptide sequences.

We strategized a taxonomic classification based on a comprehensive list of peptides obtained through ambiguation with isoBLAST and subsequent classification with ClassiCOL, an anti-Occam’s razor-inspired algorithm for decision making at each taxonomic level.

### Collagen peptide ambiguation through the novel isoBLAST approach

Inferring proteins is not the goal of ZooMS^2^ approaches. Rather, it is the species that must be inferred. To this end, we propose a different approach, applicable to any ZooMS^2^ search result output, irrespective of the instrument or search engine used.

Briefly, many search engines do not allow the user to export ambiguous results (and several spectra could potentially be matched with more than the maximum of ten peptide candidates with similar scores that are displayed in Mascot). Given that many tryptic peptides match to a multitude of different species entries in the database, the correct answer for a given spectrum is therefore frequently not exported. To maximize the chance that each spectrum contains the correct peptide-spectrum-match (PSM) amongst the list of ambiguous annotations, we developed isoBLAST.

Conventionally, when peptide sequences need to be attributed to an organism, this is done through a BLAST algorithm, which uses an evolutionary background scoring matrix to assess the similarity to protein sequences in the database. However, when looking for all the peptides in a database that could explain a given spectrum, there is no use in allowing for non-isobaric changes, since the precursor mass is accurately measured (e.g. at 10 ppm mass error) and a non-isobaric change would additionally shift the masses of the rest of the b- and y-fragment ion series to either side of the amino acid change. Therefore, isoBLAST searches the database only for isobaric changes, i.e. local, mono-to dipeptide amino changes that do not change the observed mass and are expected to have little or no impact on the scoring. As described above, these include (i) a positional isomer, (ii) isoleucine-to-leucine and N or Q deamidation (to D or E), (iii) mono-to-dipeptide changes and (iv) the isobaric -PS-to -P_ox_A- or any other isobaric changes involving a defined PTM.

Note that the initial search for each of the samples presented in this manuscript is done against the ClassiCOL database. That said, users can also provide CSV output of searches against a more constrained database like e.g. Swissprot with a given taxon filter. In case of the latter, isoBLAST will increase ambiguity using the ClassiCOL database and coordinately extend the considered organism selection beyond the constrained database used during the initial probabilistic database search. If clear evidence of unexpected organisms is found in this way, the process is best iterated on a database search performed against the ClassiCOL database.

### Taxonomic classification method

In order to tackle the species inference problem, a novel species classification algorithm was developed specifically for the ambiguated data, since this ambiguation changes the prior assumption for classification. Conventionally, 100% correct annotations are assumed, whereby tryptic peptides can be found that are unique to a given taxon or species, as is envisioned in the Unipept approach. Yet, if the search algorithm cannot distinguish between two equally scoring peptides, it will propose only one, leaving the possibility of branching off into an entirely wrong lineage of taxonomy and moving the ambiguity downstream from the search algorithm to the classification algorithm. Moreover, semi-tryptic peptides are omnipresent in paleoproteomics data due to high levels of protein taphonomy, but are still under development in the Unipept tool, which currently still results in the loss of valuable and informative peptides for species classification.

Now the axiom becomes that for each spectrum present in the data, the correct explanation is included, in an otherwise redundant list. Therefore, the algorithm (**Figure 2A**) starts by building a bi-cluster based on the available sequences. We create a taxonomic tree representing all species (according to NCBI taxonomy) matching at least 2 peptides to at least one protein sequence, i.e. no ‘single hit wonders’ are considered. Next, at every taxonomic branching point from the last vertebrate common ancestor to species, the two branches are compared one to one and can either be discarded or retained: when a branch cannot be distinguished from its counterpart because it is a subset of the other, it is discarded, yet when a difference at the peptide or protein level distinguishes it, the branch is retained (**Figure 2B**). When both branches have uniqueness to them, both are considered and the algorithm splits its search towards both branches separately, i.e. it considers the sample to be a mixture of ≥2 species. When branches cannot be further separated based on their peptide (and thus protein) content, (i.e. the peptides within both branches are identical) the algorithm halts at the taxonomic level of that branching point (**Figure 2A**). Notably, after every split, all initial peptides are reconsidered, having the benefit of retaining peptide sequences that originated from independent mutations that occurred after the speciation event (**Figure 2B**). In Unipept-like classifications, such peptides are usually plotted to the lowest common ancestor of the two species that express it, while all other species in that branch might not have this sequence. Because of the repeated decision making at every taxonomic branch, these peptides are rescued and have proven to contribute to the taxonomic classification.

**Figure 2.**
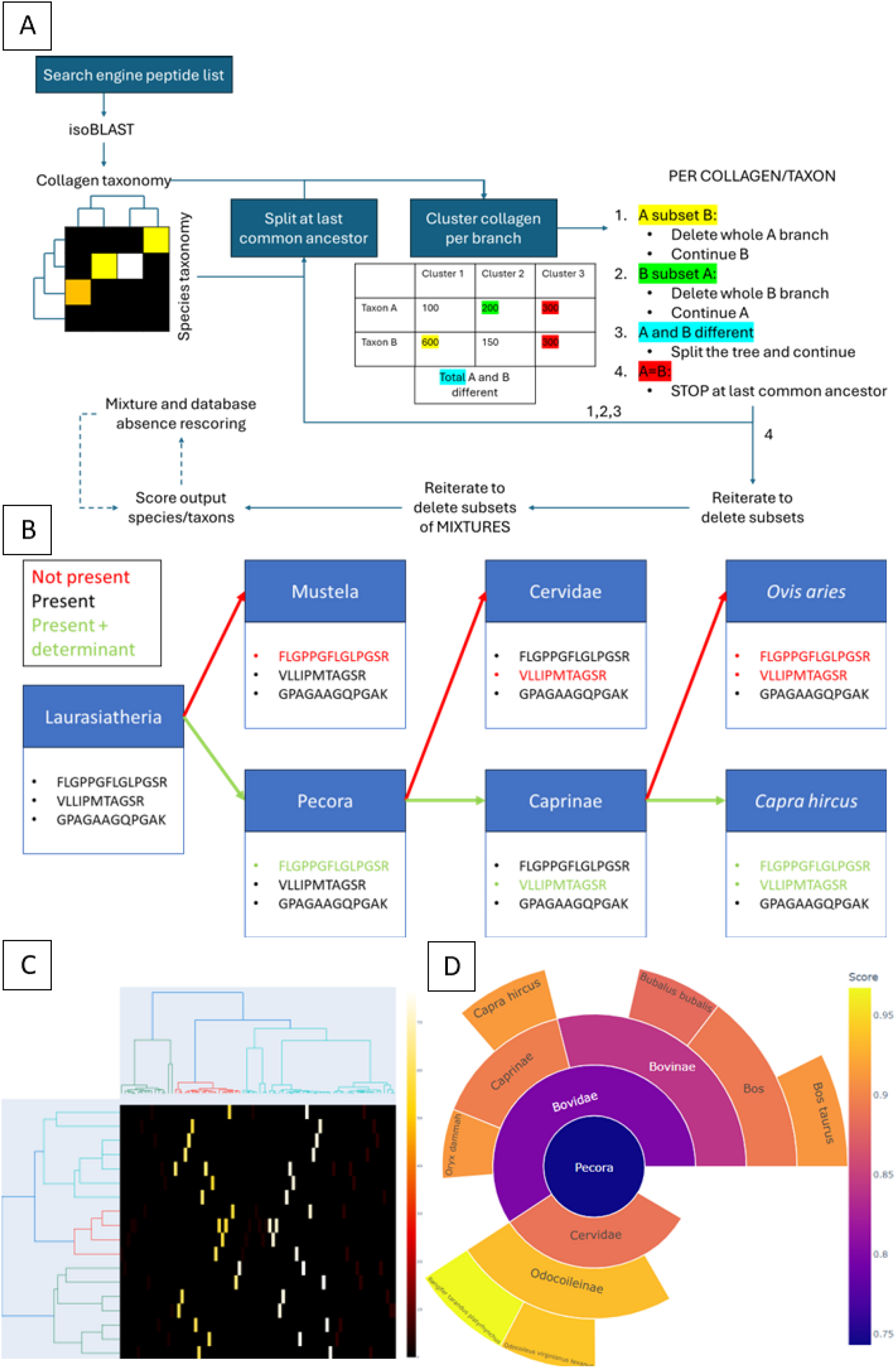
Overview of the ClassiCOL pipeline. **A**. Schematic of the ClassiCOL algorithm workflow starting from a search engine output file, the numbers represented in the table are randomly chosen to highlight the possible difference between two taxonomic branches. **B**. Schematic overview of peptide reuse from independent mutations after the speciation event, which are important for classification over different taxonomic levels. The peptides represented in the scheme are true peptides, from different spectra, that were found in the goat reference dataset from Rüther et al., 2022 (*8*) **C**. Visual representation of the taxonomic-collagen bi-cluster. The cluster is zoomed in at the Pecora family level on a sample from *Rangifer tarandus* (RMC42); the lighter the color in the heatmap, the greater the coverage for that collagen sequence. The y-axis represents the species taxonomic tree, the x-axis represents the homology between collagen sequences, colored by relatedness. **D**. Output sunburst plot of the *Rangifer tarandus* sample, where higher scores represent higher likelihood that the named taxon approximates the sample content. All ambiguity is retained in the final output, whether originating from the isoBLAST approach or directly from the search engine results. The interactive version can be found as **Supplementary Figure 2**.

**Figure 2C** visualizes this approach with a simplified heatmap, exported from the ClassiCOL tool during every analysis. On the x-axis, all considered collagen sequences in the result file are clustered through a sequence homology matrix (“Collagen taxonomy”). On the y-axis, we depict the taxonomic tree representing all species from NCBI Taxonomy matching at least 2 peptides to at least one protein sequence. It is through this matrix that the anti-Occam’s razor-inspired algorithm finds the species or taxonomic level that can best explain the comprehensive peptide-to-spectrum match space created through isoBLAST.

As for any classification algorithm, a single unique peptide difference is sufficient to retain a species. Therefore, multiple taxonomic levels and different species will be retained in the final output, reflecting the ambiguity provided by the Mascot search and isoBLAST algorithms. To facilitate user interpretation, an adapted Bray-Curtis pairwise distance dissimilarity (*41*) score is calculated by comparing the possible peptides of each species to the entire set of collagen related peptides in the sample.

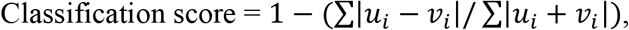

where *u* is a binary vector showing presence-absence of peptides for each species, and *v* representing a vector of all collagen related peptides in the sample. In other words, it computes the distance between the species array and the sample array, adapted to 1-Bray-Curtis distance to calculate the similarity. Thus, the more a species contributes to the sample proportionally, the more likely it is for the sample to have originated from said species. Finally, an interactive sunburst plot is provided to the user as an output that still captures the underlying ambiguity, yet is color-coded according to the score to facilitate interpretation (**Figure 2D and Supplementary Figure 2**).

When applying ClassiCOL, **three outcome scenarios** are possible (see **Figure 3**): **(i)** one single species was ascribed a higher score than all others, which includes a true species match or a single species match to the closest related extant species in the database; **(ii)** the algorithm displays the lowest non-species level taxonomy where lower branches cannot be further separated based on their peptide content, e.g. at the genus level, indicating that any member of that genus is considered to be an equally likely outcome; **(iii)** several different species are deemed to be similarly likely outcomes. In this third case, two separate explanations can be offered, depending on the relatedness of the species. First, when the two species are distantly related, this reflects the output of a physical mixture of species present in the sample, which may also derive from contamination. Second, both species are very closely related, yet explicitly depicted as having evidence for being in the sample (as opposed to the algorithm stopping the bifurcation at a higher taxonomic level as in (ii)). This happens when the species to which the sample belongs is not represented in the ClassiCOL database, i.e. either because the sample species is extinct, or the sample is from an extant species which is not in the protein database, like for example a roe deer (*Capreolus capreolus*). Irrespectively, we consider these to be “*genetic mixtures”* in the perspective of the database used.

**Figure 3.**
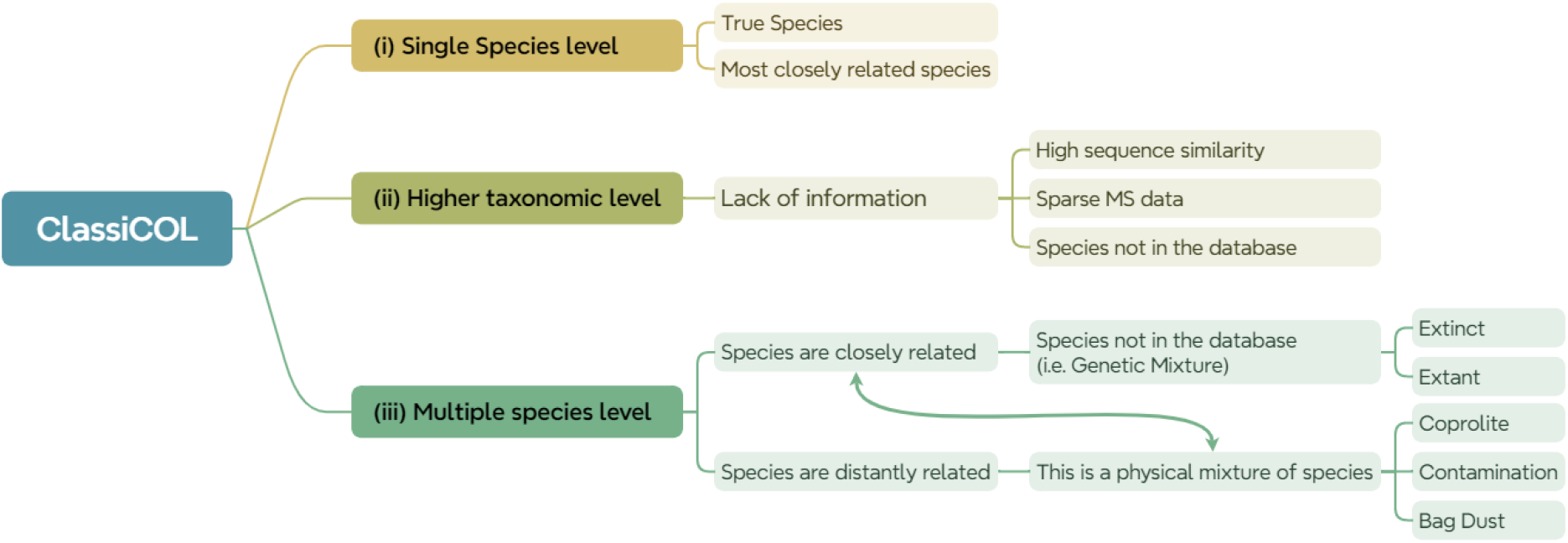
Overview of the ClassiCOL outcome scenarios. This schematic (created with xmind) shows all potential outcome scenarios of the ClassiCOL algorithm in the form of a decision tree. (i) Depicts the scenario of the identification of a single species, which can be the true species or the most closely related species in the database, (ii) shows how to interpret an outcome specific to a higher taxonomic level than the species level, and (iii) visualizes how to infer a physical and/or a genetic mixture from the ClassiCOL output files.

Users should note that the classification algorithm has been programmed with the purpose of species identification based on all collagen isoforms, as these proteins have been known to persist over the longest periods of time and in high abundances. However, the algorithm does not necessarily rely on collagens alone, opening up the possible addition of non-collagenous proteins (NCPs) to the database in the future. Alternatively, a non-collagen custom database could be used. In fact, NCPs are typically less evolutionarily conserved and can be annotated more efficiently with an order-, family-, genus- or species-specific database after the bone sample has been classified to said level on the basis of collagen content (*42*), although we do not extend on this strategy here. In this context, it is also relevant to note that the computational time increases linearly with both the number of peptides in the result file and the number of considered species in the ClassiCOL database, emphasizing the crucial role of bioarcheological preassessment. In the event that the user deploys prior knowledge to restrict the taxonomy outcome of a sample (e.g. to Mammals), the algorithm will still consider a maximum of 15 species from each of the Vertebrate Classes that do not belong to that taxonomy (e.g. vertebrates other than Mammals) as entrapment validation sequences, i.e. known false targets (*43*). If the user chooses to restrict the results to species within the Pecora infraorder, the algorithm will still choose other non-Pecora members of the Mammal vertebrate class as entrapment targets, which maintains the possibility of false targets both closely and distantly related to the sample species.

### Algorithm performance on public datasets

Different LC-MS/MS techniques and search algorithms have been reported for species identification using ZooMS^2^ in recent years, all using a different (user-driven) decision making process, including - most recently – MS^3^ spectra (*39*). Therefore, to increase user-friendliness to all users, regardless of expertise level, the Collagen Classification (ClassiCOL) algorithm was developed as a common post-processing pipeline that requires only a list of identified peptides as input, including PTMs and their respective localization. In order to demonstrate its potential and applicability towards a variety of different LC-MS/MS acquisition methods and sample types, we processed several publicly available datasets. In the process, we demonstrate how the different outcome scenarios can be interpreted (**Figure 3**). All individual interactive sunburst plots and an overview table of all the scorings for the public datasets are compiled as **Supplementary data 2-3**. As shown in **figure 4**, our annotations agree well with the published interpretations.

**Figure 4.**
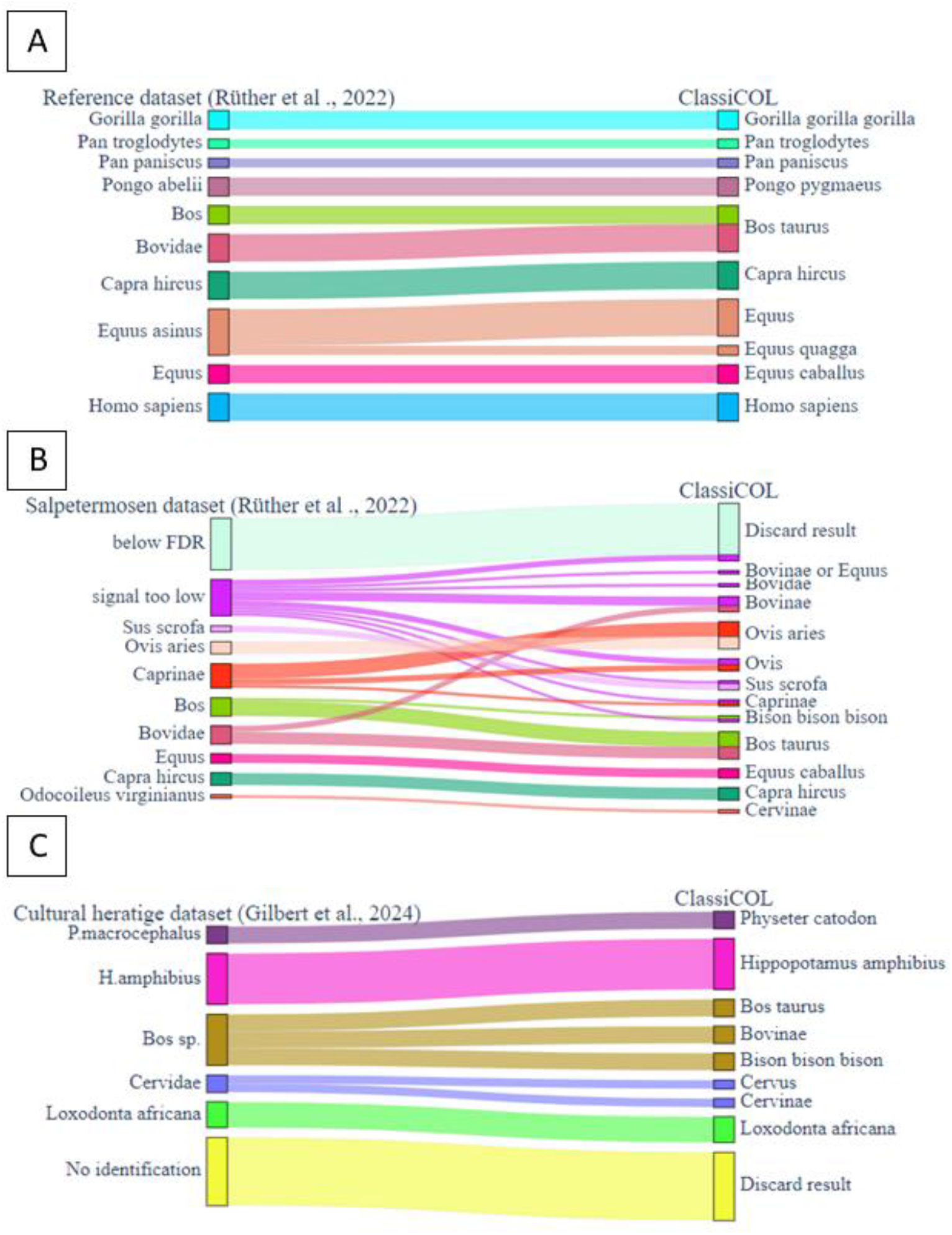
Similar, yet finer grained classifications are computed by the ClassiCOL pipeline compared to the original papers’ proteomics results. **S**ankey diagrams show the taxonomic classification approximation made by the ClassiCOL pipeline in comparison to the proteomics outcome of **A**. the reference collection and **B**. the Salpetermosen datasets from Rüther et al, 2022 (*8*), and **C**. a cultural heritage dataset from Gilbert et al., 2024 (*39*) Colors are specific to the classification of the original paper per taxonomic level, and are directed to the classification made by the ClassiCOL algorithm. We observe that ClassiCOL can also rescue outcomes from samples with poor data quality, as seen for the (B) Salpetermosen ‘signal too low’ input.

First, we demonstrate the performance of the tool on a reference dataset produced by Rüther et al., 2024 (*8*) (**Figure 4A)**. Briefly, they describe a workflow wherein they made a site-specific difference matrix, only considering single amino acid mutation sites that differ between two species. This matrix was then used to score LC-MS/MS data after it was mapped to a multiple protein sequence alignment from multiple species in their database. They combined this metric with peptide intensities, peptide counts and precursor counts, giving more weight to higher quality MSMS spectra to overcome ambiguity. Our results concur with their original proteomic identifications; both are in line with the reference collection bone morphologies. Still, we identified one species differently as being *Pongo pygmaeus*, which matches the morphological identification and not the *Pongo abelii* identification from Rüther et al.’s LC-MS/MS workflow (**Supplementary Figure 3**). In this case, this derives from the fact that the correct protein sequence was not present in the database used in the original paper. On the other hand, we identified one of the reference samples as originating from *Equus quagga* rather than *Equus asinus*, the latter of which was morphologically determined (**Supplementary Figure 4A**). In this specific case, the recycling of peptides at each taxonomic decision level (**Figure 2B**) resulted in a high score given to a single peptide specific to zebra and horse but not to donkey, while all other peptides retrieved from the donkey sample are shared between zebra and donkey, therefore zebra was a more likely outcome (**Supplementary Figure 4B**). Lastly, we managed to find evidence for three equine species when analyzing two mules in the reference collection dataset, which corresponds to their mixed parentage from both *Equus caballus* and *Equus asinus* (**Figure 3ii, Supplementary Figure 5**). This particular form of hybrid genetic relatedness can be further interrogated through peptide-species match overlaps, as discussed later.

Next, we reanalyzed the samples excavated at the Salpetermosen site in Denmark (*8*) (**Figure 4B**). Because a lot of archeological samples yield little high-quality data, several of these samples were classified as ‘signal too low’ in the original manuscript. However, with our approach, leveraging the increased ambiguity that inevitably results from the low-quality data, we were able to classify several additional samples which all matched the morphological species classification (**Figure 4B, Supplementary data 2-3**). Furthermore, samples that were previously classified to the subfamily level (e.g. Caprinae), are now classified to the species level, again matching the morphology and demonstrating the performance of the algorithm (**Supplementary Figure 6**). Additionally, one sample that was initially classified as *Bos* returned a likely match to a kind of bison, which could be derived from a species not in our database, e.g. *Bison bonasus*, given the age of the sample and the geographical location of the excavation site (**Supplementary Figure 7**).

Lastly, there is also a lot of interest in identifying paint binders and bone- or hide glue in cultural heritage collections (*44*–*47*). Therefore, we tested the applicability of the workflow in cultural heritage science, first focusing on data from a paper that identified the species of origin from bone and ivory museum pieces from the Smithsonian Institution, produced by Gilbert et al., 2024 (*39*). (**Figure 4C**). Here we were able to identify the species of origin without going into each spectrum separately and consulting the MS3 spectra created by fragmenting fragments from the MS/MS spectrum, as was necessitated by the original approach. Furthermore, we were able to pinpoint some samples to a more specific taxonomic level compared to the original manuscript. In fact, one sample was identified to have originated from a species of Bison, which was not considered in the paper (**Supplementary Figure 8**). Also, as stated in the original paper, one of the artefacts was covered in glue, explaining the *Bos taurus* identification of an ivory object (*39*). This shows that our approach can also classify species based on collagen types other than I and II, such as COL3A1, which is a marker protein for hide-based glues which are commonly investigated in heritage paleoproteomics (*47*) (**Supplementary Figure 9**).

We refer to **Supplementary Data 3** for a full overview of this validation on public data. Overall, incorrect species classification and annotation to an irrelevantly high taxonomic level were almost exclusively found in samples displaying a score below 0.6 for a single species and/or with fewer than 20 unique collagen peptides (peptidoforms). These criteria were therefore used as fixed thresholds to avoid ambiguous classification.

### Classification of a wider selection of in-house processed samples

To extend on the taxonomic coverage, we extracted proteins from extinct and ancient species from Belgian archaeological sites and acquired the data through LC-MS/MS (depicted in **Figure 9**). First, a number of Ahrensburgian, Mesolithic and Neolithic samples were selected from the Belgian archaeological sites of Remouchamps (*n*=1) (*48*), L’Abri du Pape (*n*=14) (*49*), and Oudenaarde Donk (Neo 1) (*n*=41) (*50*), including fish remains and microfauna. Similarly to the validation on public data, almost all sample results matched the initial morphological classifications, yet with greater taxonomic depth and with a few exceptions (see red arrows in **Figure 5**). In particular, one sample that was morphologically classified as *Cervus elaphus* (ID: ADP006) came out as *Sus scrofa* (**Supplementary Figure 10**). Morphological reinspection of the proximal epiphysis revealed that it was indeed an exceptionally large wild boar specimen. Another example involved a bone originally classified as fox (*Vulpes vulpes*, ID: ADP0014), which was classified by ClassiCOL as otter (*Lutra lutra*), an identification which was again confirmed through secondary morphological classification (**Supplementary Figure 11**). Conversely, we also identified a sample as red deer which was morphologically determined to be roe deer (ID: ADP0011), but in this case the thickness of the cortical bone excluded both adult and juvenile red deer (**Supplementary Figure 12**). Thus, these very closely related species of cervids could not be disentangled by the algorithm, which can be largely accredited to the absence of the *Capreolus capreolus* in the database and highly similar collagen sequences within the family Cervidae, an issue aggravated by the poor coverage of collagen peptides in this sample.

**Figure 5.**
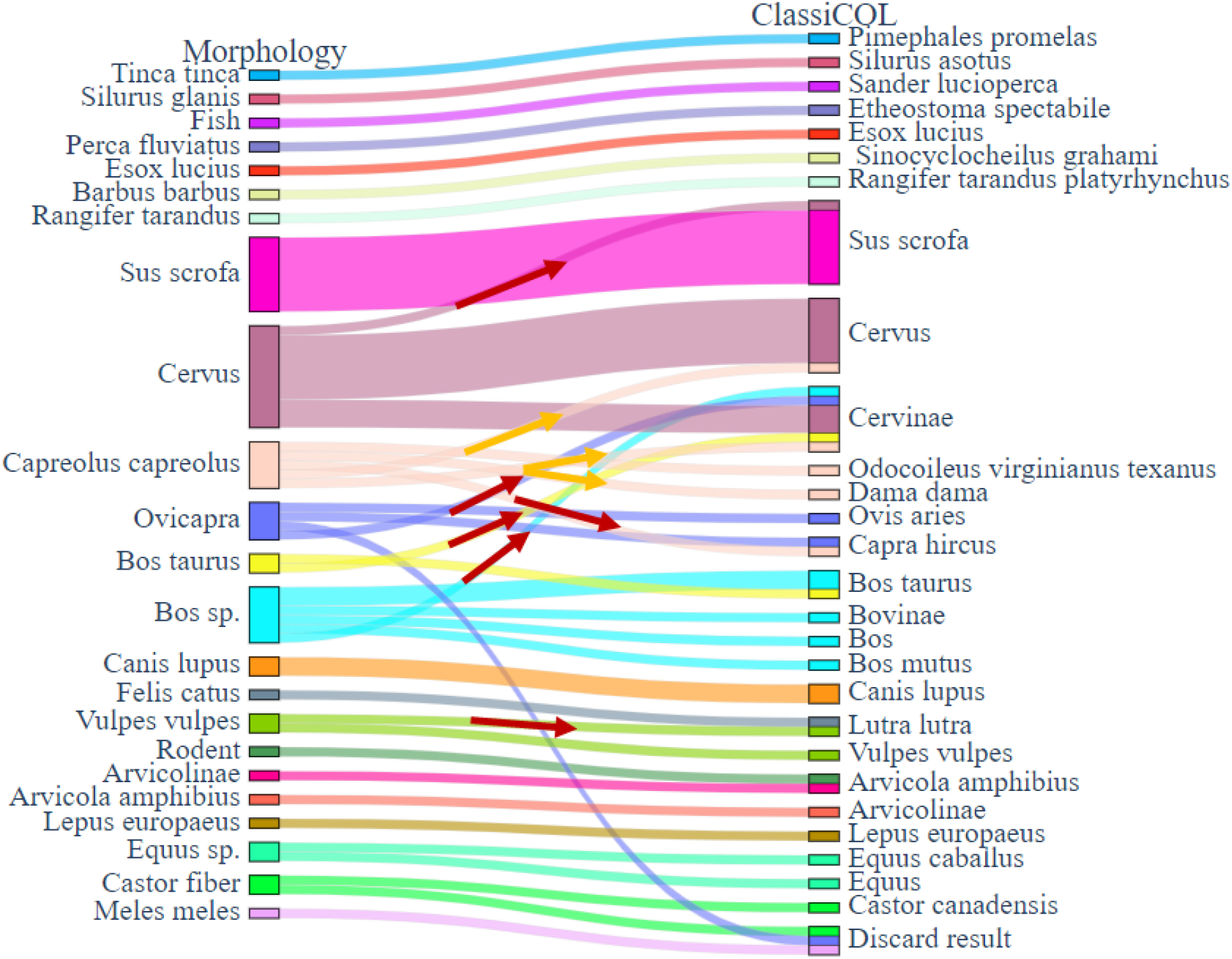
Analysis via ClassiCOL showed a few dissimilarities with bone morphology for in-house processed samples. Connection graph shows the classification made by ClassiCOL on samples from ADP, RMC and OD. Red arrows indicate changes of classification to distantly related taxa based on the proteomics data. Yellow arrows show discrepancies with the morphological estimation within the same family.

Some classification discrepancies were impossible to morphologically reassess as the bone fragments were too small, too degraded, or too similar to closely related species to give an accurate re-assessment (IDs: OD024, OD08, OD042, OD029 and OD31). (**Supplementary Figure 13-17**) Regarding the fish fragments, we saw that several of the bones that were morphologically identified as fish were indeed classified as fish, yet often to a higher taxonomic level (**scenario Figure 3ii**), incentivizing us to expand the database to 168 species of ‘fish’ (including Actinopterygians and Chondrichthyans). The same analysis was performed, and gave a more accurate species approximation compared to when the species of origin was not considered in the first search (**scenario Figure 3i)**. Indeed, all actinopterygian specimens were identified to have originated from the morphologically estimated genus or species, when included in the database (**Supplementary Figure 18-23**). Notably, several fish genera are not yet present in any database. These species returned results indicating a mixture of close relatives under the same taxonomic family. Next, we assessed the performance of the algorithm on actual mixtures and on species not included in the database, including extinct animals.

### Challenging samples: Rescoring mixtures and unsequenced species

With the tool’s performance well-established on bones that were sampled specifically for the identification of species, we turned to applications in paleoproteomics that require the identification of different species from the same sample (**Figure 3iii)**. The ability of the ClassiCOL approach to retain multiple species outputs per sample, coupled with the scoring approach, enables inference of physical as well as genetic mixtures, including extinct species.

First, to better resolve cases of extant and extinct species not present in the database, we strategized a data-driven selection of the top five candidate taxa represented in the sunburst output, followed by a recalculation of the Bray-Curtis score on the discriminant peptides only, i.e. a second-pass search. From the in-house-processed fish remains, we isolated a one-species and a multiple-species sunburst (**Figure 6A and 6D**), isolated the unique peptides (**Figure 6B and 6F**) and recalculated the Bray-Curtis score (**Figure 6C and 6F**) in a second-pass classification. For a single high scoring species, the second-pass rescoring can still offer a higher score drop-off to increase the certainty of the annotation (**Figure 6C**). Yet, in the case that a species is not in the ClassiCOL database, this drop-off is less pronounced, i.e. a gradual decline in scoring is pertained, approaching a more detailed relatedness of a genetic mixture. Thus, while not obligatory for single species, this extra algorithmic step leads to a higher resolution in classification, much like second-pass Percolator rescoring can improve peptide annotation in database searches (*29*), especially when dealing with physical and/or genetic mixtures.

**Figure 6.**
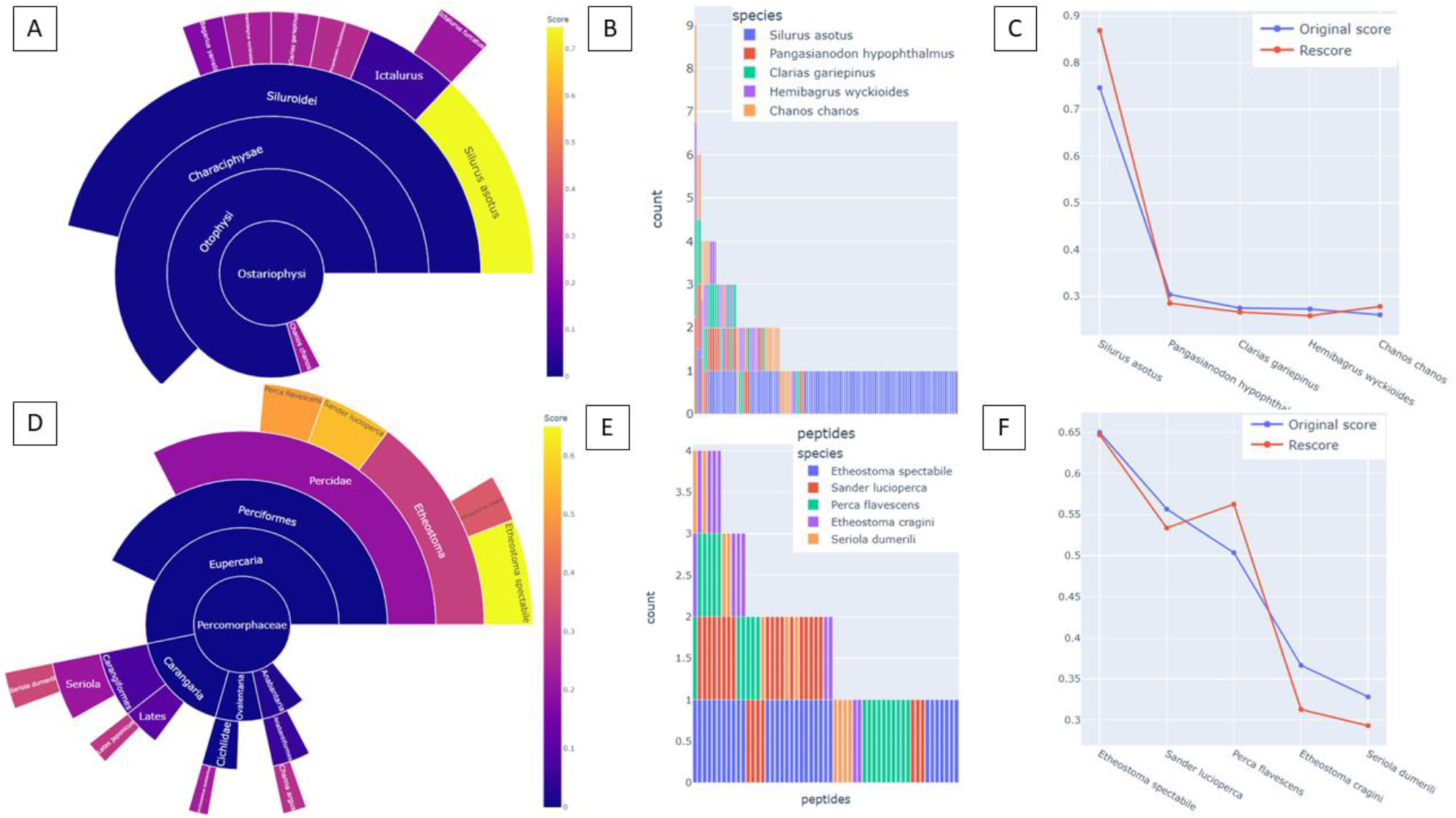
Rescoring highlights instances of genetic relatedness when the true species is not in the database. **A-C**. A fish bone morphologically identified to the genus *Silurus* (OD0004) analyzed with ClassiCOL. The barplot (**B**) and line graph (**C**) visualize high uniqueness for *Silurus asotus* both before and after re-scoring, with a large drop-off for other species scores. The species *Silurus asotus* is likely the closest related species in the database to the species of origin - *Silurus glanis*, based on the geographical location and time period. **D-F**. Rescoring of morphologically estimated *Perca fluviatus* (OD0002) with ClassiCOL shows potential for genetic relatedness with other members of the family Percidae. Barplot (**E**) shows a significant non-overlapping difference between sample peptides shared with *Etheostoma specabile, Sander lucioperca* and *Perca flavescens*. Line graph (**C**) demonstrates similar re-scoring distributions, suggesting that the true species is not included in the fish database, but can be found in the family Percidae.

Next, we assessed how this approach reflects extinct species. Therefore, we reanalyzed a public dataset from Bray et al., 2023 (*51*), which was focused on >100,000-year-old remains excavated in Scladina Cave in Belgium (**Figure 7A**). Here as well, all classifications matched the taxonomic level presented in the manuscript, yet most samples were classified to the species level present in the database, i.e. an extant relative. Strikingly, several samples resulted in a sunburst plot that looked more like a mixture. As an example, a cave bear bone resulted in an ambiguous annotation of ursine species, which is not the same as the algorithm stalling at a higher taxonomic level. Still, this shows that extinct animals that are not in the database will be challenging to interpret, mimicking sample mixtures, and giving ambiguous results of two closely related species at best.

**Figure 7.**
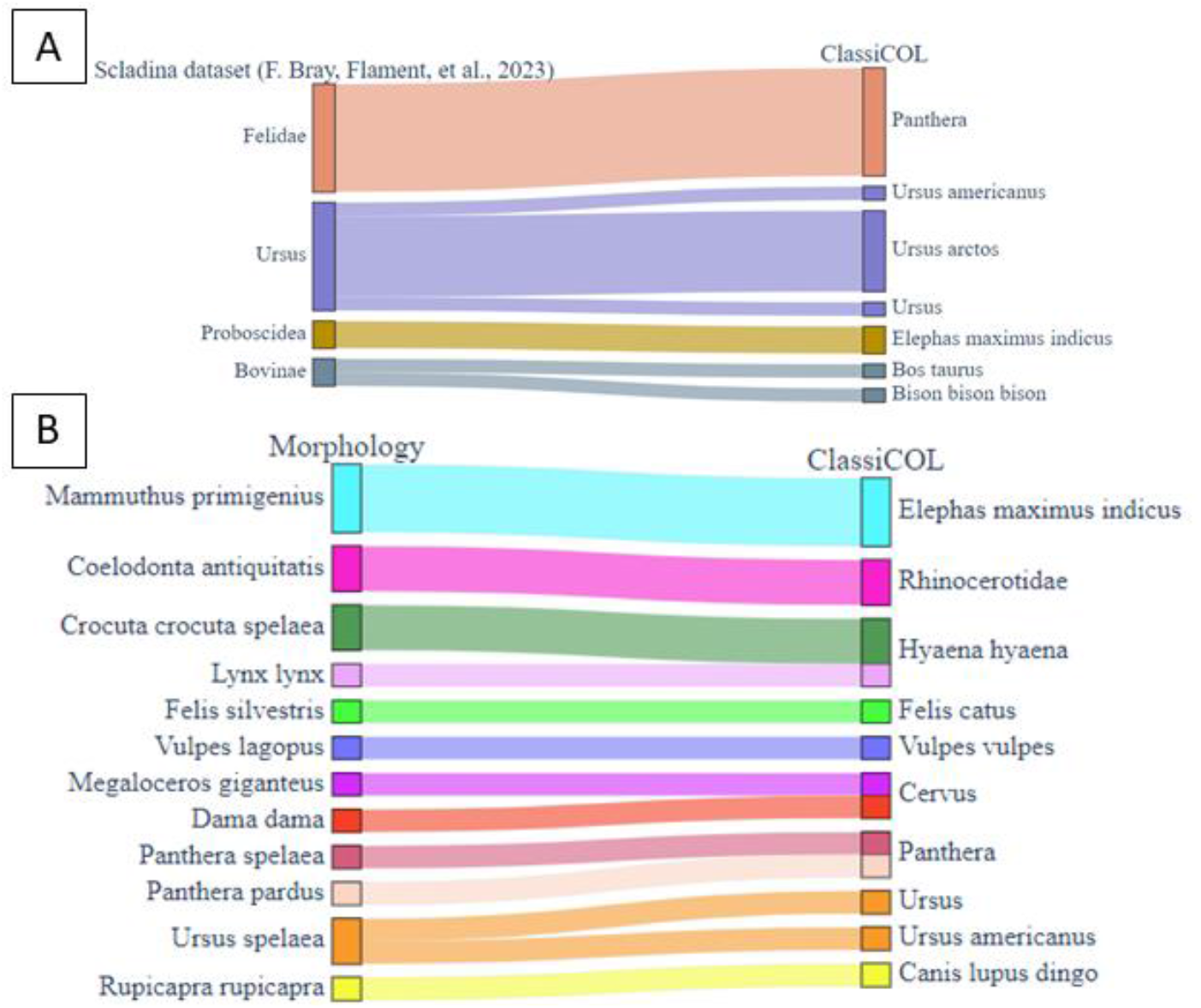
Performance of ClassiCOL on extinct mammals. **A**. Reanalysis on the publicly available data of Bray et al., 2023 (*51*), provides identifications matching the original publication, with taxa identification where manual curation was needed in the original paper. **B**. ClassiCOL’s performance on in-house processed extinct mammal samples excavated from the same site as Bray et al., 2023, being Scladina Cave (*51*) both sample bag dust as bone dust was analyzed.

In parallel to the Ahrensburgian, Mesolithic and Neolithic samples, we processed and analyzed several extinct animal specimens from Scladina Cave, the same site as Bray et al., 2023 (*51*). Again, we observed that the species classification agreed with the morphological classification (**Figure 7B**). Here, we were able to classify *Coelodonta antiquitatis* only to the family Rhinocerotidae because of the absence of woolly rhino and the closest living relative in the database (**Supplementary Figure 24**). However, due to the geographical location of the site and Upper Pleistocene dating of the bone, woolly rhino was the only possible outcome. Again, this emphasizes that interdisciplinary expertise will always be required for robust classification.

Additionally, two samples had to be morphologically reassessed due to discrepancies with the ClassiCOL output. The samples formerly identified as being *Lynx lynx* and *Rupicapra rupicapra* were classified by ClassiCOL as *Hyaena hyaena* and *Canis lupus dingo* respectively (**Supplementary Figure 25-26**). The *Hyaena hyaena* was confirmed as the mandible most likely originated from a cave hyena cub and the *Canis* identification was confirmed due to shape, curvature, and muscle attachments on the bone of the juvenile specimen. For this analysis, when possible, we sampled the bone both directly and indirectly as disintegrated bone dust in the storage bag, each of which produced highly covered collagen sequences (COL1A1 and COL1A2) and taxonomically classified the sample correctly according to the morphology. In other words, both direct invasive sampling and swabbed bone dust from containers can be used for analysis. Thus, limiting destructive sampling is an exciting development as minimally destructive sampling is important for responsible heritage curation and paleoproteomics as a research field (*52*).

Physical mixtures are of great interest, e.g. because bags or boxes commonly contain several bones that can derive from different species, which could theoretically all be identified in a single run. Another challenging prospect is the analysis of coprolites, which can contain mixtures of extinct animals. Therefore, we analyzed the dust from a bag containing cave hyena coprolites (*Crocuta crocuta spelaea*) excavated in Scladina Cave (Layer T-RO, early Aurignacian). These paleofaeces preserve information on carnivore meals and potentially collagen bone fragments from multiple extinct animals, as hyena are opportunistic scavengers. **Figure 8A** shows the sunburst plot depicting distantly related species that point towards a physical mixture and closely related species within each of these branches, pointing to a genetic mixture, in turn potentially indicative of extinct species. During mixture re-scoring, we therefore allowed the algorithm to take more than the default five and thus to include all possible outcome species (**Figure 8B**). Via the aforementioned re-scoring, the potential candidates in the mixture were confirmed (**Figure 8C**): all three of its replicates contained two high-scoring distantly related species: European Bison and Cave bear (**Figure 8**).

**Figure 8.**
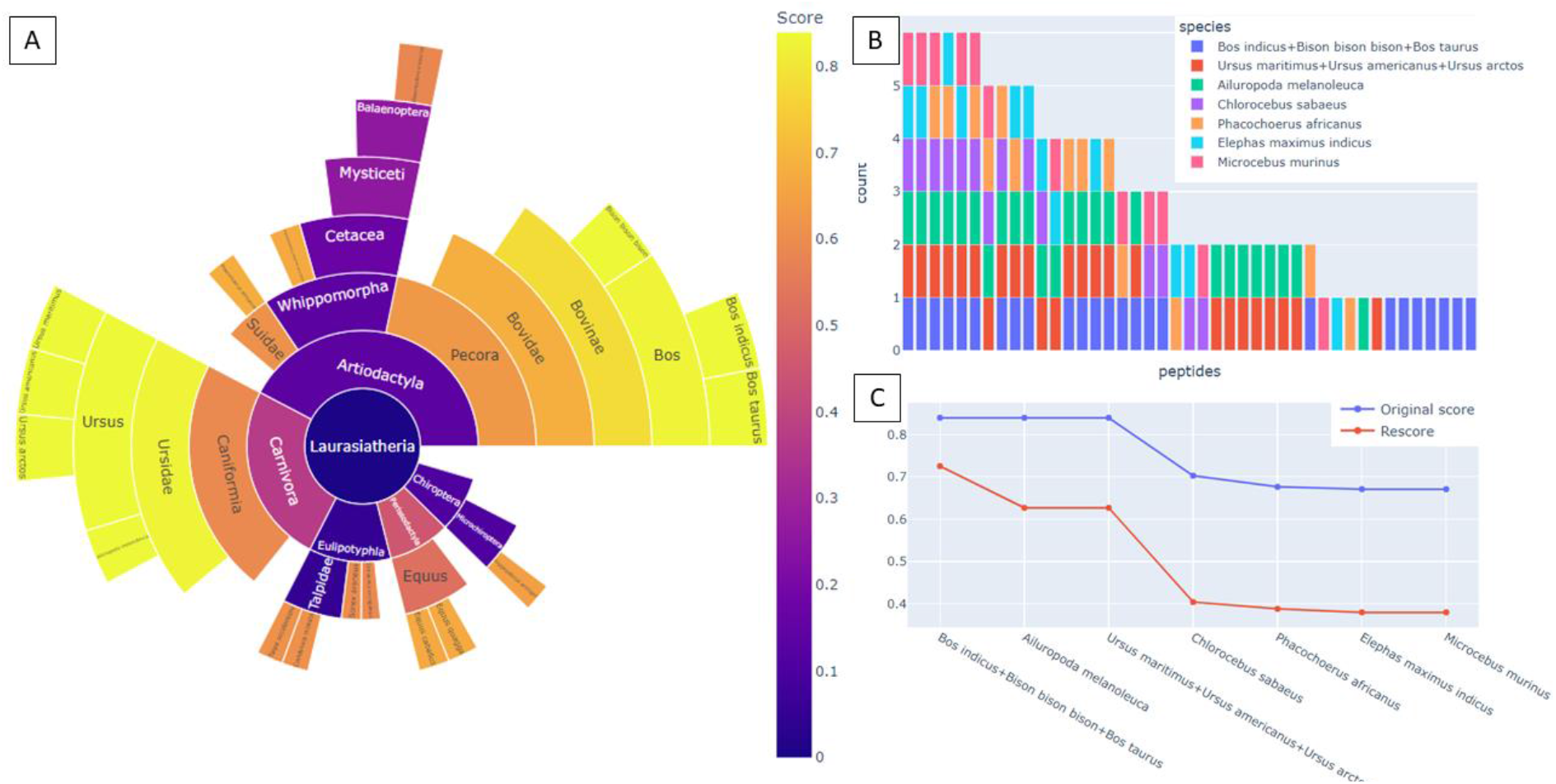
Analysis of a cave hyena coprolite via ClassiCOL with subsequent mixture reanalysis. **A**. Output of ClassiCOL shows high scores for two distantly related species, indicating a sample mixture. **B**. Barplot depicts 1 isoBLAST peptide difference amongst bears and high uniqueness for both Ursus as Bovinae. **C**. Rescoring of the showed no drop-off between bear groups nor was a drop-off seen for Bovinae, a drop-off is noticeable after Ursus. These results show that the sample is comprised of both Ursine and bovine bone fragments.

## Discussion

Our novel ClassiCOL pipeline has the ability to approximate or identify the species of origin for archaeological bone samples starting from the search results of any LC-MS/MS run (ZooMS^2^). These analyses are intrinsically more information-rich than a ZooMS analysis because they contain peptide fragment data, yet the ambiguity in search results has long hampered their routine implementation. Therefore, we here embrace and extend the ambiguity found in searches against an extensive manually curated ClassiCOL collagen database, using our isoBLAST approach, to ensure that the correct annotation for each spectrum is always present. With this new premise, the ClassiCOL taxonomic classifier is then applied to isolate the species that explains most of the potential peptide candidates in the isoBLASTed result space. By comparing the results of our algorithm to several publicly available datasets, we have demonstrated that the ClassiCOL algorithm matches existing results, routinely gives more precise species identifications and can rescue poor quality data. In contrast to other approaches, ClassiCOL is not reliant on unique peptide stretches, but can differentiate between species based on overall peptide content, provided through a search engine output file in CSV format, including the reconsideration of peptide stretches representing independent mutations after speciation events. As the collagen database consists of all collagen types, heritage samples containing bone or hide glue can also be analyzed through the same pipeline without any adaptations. Finally, our approach shows promise to resolve genetic mixtures, i.e. species not in the database, whether extant or extinct, as well as different species in a physical mixture, as demonstrated on dust from coprolites.

The main limitation of ClassiCOL is the interpretability of the final sunburst plots. Therefore, we provide examples of the three main scenarios of outcomes and guide the user by including a decision tree. Still, the very fact that all potential peptide candidates need to be considered during classification, in combination with sample complexity and degradation, sometimes leads to complex sunburst plots that require thorough curation. In the end however, ClassiCOL is an analytical algorithm developed only to guide decision making and all analyses should be pre- and post-assessed by a zooarcheologist for contextualization.

## Methods

### The Archeological Contexts of Oudenaarde-Donk (OD), Abri du Pape (ADP) and Remouchamps (RMC)

A single *Rangifer tarandus* element from Grotte de Remouchamps was sampled. Located in Southern Belgium within the province of Liège near the Amblève River (**Figure 9**), the cave has been known since the 18th century and attracted the attention of various prospectors. E. Rahir (*53*) and later M. Dewez (*54*) conducted the latest and most complete research on the archeological material uncovered during their excavations. The material, lithic, and faunal remains are associated with Ahrensburgian occupation(s) dated from ca. 12,700/12,500 to ca. 11,400/11,200 cal BP, corresponding to most of the Younger Dryas and the early Preboreal (*48*).

**Figure 9.**
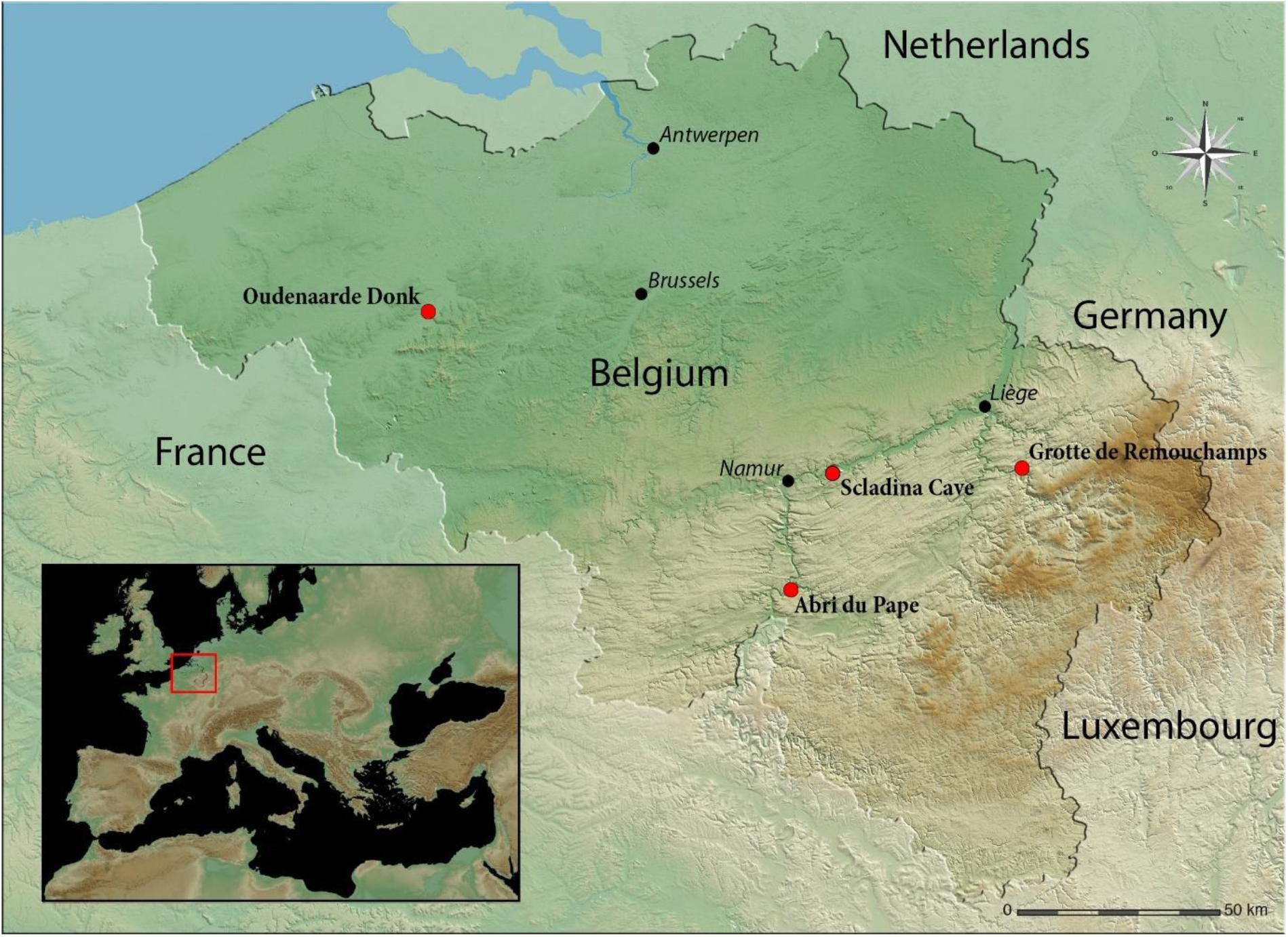
Geographical location of the Belgian archeological sites. Oudernaarde-Donk (OD), Scladina cave, Grotte de Remouchamps (RMC) and Abri du Pape (ADP) are represented by red dots. Map generated with global mapper.

The rock shelter of Abri du Pape is located in the province of Namur, in southern Belgium, on the right bank of the Meuse River (**Figure 9**). Initial excavations at the site were conducted by Ph. Lacroix in 1988, who uncovered archaeological deposits in a sondage. This early work was followed by a series of major excavations throughout the 1990s, led by J.M. Leotard, M. Otte, L.G. Straus and a multidisciplinary team (*49, 55*–*57*). The site revealed an extensive sedimentary sequence documenting human occupations spanning from the Mesolithic to the present day. The 14 faunal remains sampled for our analyses come from the Mesolithic layers, dated from ca. 7968 to ca. 6509 BCE, and were initially analyzed by A. Gautier (*49*).

Finally, the 42 faunal samples from Oudenaarde-Donk were excavated from Neo 1, one of the 10 sites identified at this location on the left bank of the Middle Scheldt (**Figure 9**) (*58, 59*). Several faunal remains, among human remains (not analyzed here), were directly radiocarbon-dated. The C14 dates of the faunal remains range from 6177 to 3178 cal BP, indicating a Middle Neolithic to Early Bronze Age origin (*50, 60*).

Bone powder was obtained from the exterior surface of the bones during cleaning prior to collagen extraction for stable isotope analysis. Drilling was undertaken with diamond-tipped drill bits which were changed and cleaned between each use. Cleaning was twice performed by 5 min sonication in Milli-Q water followed by 5 min sonication in 70% ethanol. A maximum of 5 mg of powder was separated from the cleaning layers and taken forward for protein extraction.

### Sample information Scladina Cave

Scladina Cave is located on the right bank of the Meuse Valley, between Andenne and Namur (**Figure 9**). The cave was discovered by amateur archaeologists in 1971 and has been under a permanent scientific archaeological programme since the early 80’s (*61*). The cave yields numerous archeological occupations mostly by Neanderthals, as well as, some evidence of the early anatomically modern humans in north-western Europe (*62*). The stratigraphic sequence is composed of no less than 120 layers for an approximate cumulated thickness of around 15 meters, which covers the Holocene up to at least the Late Middle Pleistocene (*63, 64*). Although the site is known for its archeological occupations, the sediments at Scladina Cave have preserved an impressive quantity of bone and dental remains, mainly belonging to cave bears (*Ursus spelaeus*), enabling to trace their evolution and adaptation in well-controlled stratigraphic contexts (*65, 66*). Among these cave bear remains, several bone fragments were used as retouchers by Neanderthals, documenting specific interactions between Neanderthals and carnivores (*67, 68*).

The samples from Scladina Cave were excavated from a variety of stratigraphic layers (*64, 69*). Tooth dentine from *Mammuthus primigenius* (SC1997-150-1) originated from layer Z1, as did the mandible of the *Crocuta crocuta spelaea*; for both specimens, dust from within the container was sampled in addition to the dentine/bone fragments themselves. The *Coelodonta antiquitatis* bone (SC1995-279-475) excavated from Unit 6 was sampled in a similar manner. Additionally, bone dust samples were collected from *Rupicapra rupicapra* (SC1984-543-3) and *Dama dama* (SC1986-1270-224) excavated from Unit 5. Bone dust was also collected from layers V grise - Vb of *Panthera pardus* (SC1982-284-1). Bone samples from Units 4 included *Felis sylvestris* (SC1983-57-33, Unit 4A) and *Alopex lagopus* (SC1983-80-2, Unit IV). Finally, dentine from a *Megaloceros giganteus* (SC2011-210-1, Layer 1B-RA) tooth specimen from layer 1B-RA and bone dust from *Lynx lynx* (SC2002-699-5, Layer 39) were sampled.

The plastic bag containing the coprolite samples was sampled by swabbing the interior of the bag with a sterile swab. The coprolites were excavated from layer T-RO that has yielded evidence for an Aurignacian occupation dated between 40,150–37,500 cal BP (*62*).

### Protein Extraction

Each sample was demineralized in 600µl of 0.6M HCl (Chemlab, CL05.0312.1000) for 24h at 25°C, while shaking at 750rpm (Eppendorf Thermomixer comfort). The samples were pelletized via centrifugation after which the supernatant fraction was removed and stored at -20°C as a back-up. The pellet was washed with ice-cold acetone (Sigma-Aldrich, 179124-1L) and resuspended in extraction buffer (5% SDS (Invitrogen, 15553-027) + 50mM TEAB (Sigma-Aldrich, 90360-100ML)). DTT (Chemlab, CL00.0481.0025) was added to a final concentration of 20.8mM to reduce the disulfide bridges for 30min at 37°C in the dark. Next, MMTS (Sigma-Aldrich, 64306-10ML) was added to a final concentration of 20mM to alkylate the sulfide groups for 10min at room temperature in the dark. The denatured proteins were precipitated with phosphoric acid (Chemlab, CL00.0605.1000) at pH 1.

Proteins were trapped on HiPure Viral Mini column (Magen Biotechnology, China; C13112) after addition of 165µl of binding/washing buffer (100mM TEAB in 90% Methanol (Chemlab, CL00.1377.1000)). The columns were centrifuged for 30 sec (4000 rpm, 25°C, Eppendorf centrifuge 5417R) between each of the three washing steps to elute the binding/washing buffer, and the first elution was reloaded on the column to decrease potential protein loss. The proteins were digested on-filter with 1µg trypsin/Lys-C (ProMega, V5073) in 40 µL of 50mM TEAB overnight at 37°C. The peptides were eluted from the column with 30µl of 50mM TEAB, followed by 30µl of 0.1% formic acid (FA) (Biosolve, 2324) and finally 30µl of 50% acetonitrile (Chemlab, CL00.0194.1000). All three elution steps were performed with a 1min incubation and centrifugation step. The samples were vacuum-dried and resuspended in 0.1% FA for LC-MS/MS analysis.

### LC-MS/MS data acquisition via ZenoTOF MS

The samples were analyzed using a Waters Acquity MClass UPLC system coupled with a Sciex ZenoTOF 7600 mass spectrometer in data-dependent acquisition mode. The peptide samples were trapped on a YMC triart C18 guard column, 3µm, 5×0.3mm and separated on a YMC Triart C18, 3µm, 150×0.3mm analytical column using an optimized non-linear 20 minute gradient of 1.5 to 36% solvent B (0.1% FA in acetonitrile) in solvent A (0.1% FA in water). Precursor scans (TOF-MS) were acquired for 0.1s over a mass range of 300-1600m/z. Up to 40 precursors with an intensity threshold of 150 counts per second, a dynamic exclusion of 6 seconds after 2 occurrences and a charge state between 2 and 5 were fragmented per cycle using collision induced dissociation. The fragment spectra were acquired for 0.015s over a mass range of 100-2000m/z, resulting in a cycle time of 0.920s.

### Manual curation of collagen databases

The collagen database contains a manually curated list of protein sequences collected from the UniProt and the NCBI protein repositories. Here, for each species (*n*=614), all 45 types of collagen were extracted if they were present in one of the repositories and downloaded in FASTA format (**Supplementary data 1**). Protein sequences were collected from the NCBI data repository, between the 30^th^ of October 2023 and the 28^th^ of July 2024, by blasting protein sequences from closely related species via BLASTp (NCBI). All FASTA files were combined into a single file (*n*=22,017), which was submitted on the MASCOT server. This FASTA file is a living document to which proteins are added continuously when new species are submitted into the aforementioned repositories.

### Algorithm pipeline explained

The ClassiCOL algorithm is programmed as follows:

1. As input, a Mascot CSV or a MaxQuant TXT result file is required. From the input file the peptides, protein names, modifications, location of the modifications and spectrum titles are extracted. The protein names are used to filter out any keratin, trypsin and lysC contaminants. Also, all duplicate peptides are filtered out, so each peptide is only considered once. Next, all modifications are matched with the UniMod database to extract their mass changes. With this set of considered PTMs, the “isobaric matrix” is built, containing all possible isobaric switches up to a combination of maximum 2 amino acids including the modified amino acids conform to the extracted PTMs.
2. Database selection: by default, the ClassiCOL database containing all curated collagen sequences is used, yet the user can reduce the search space to one or more taxonomic levels.
3. After extraction of all information from the search engine output file and the selected database, the isoBLAST algorithm generates a comprehensive list of potential peptide candidates. For each of the peptides three scenarios are possible:
  a. the peptide exactly matches a peptide in the ClassiCOL database.
  b. the peptide matches a sequence in the database that has ambiguity (B, Z, X amino acid annotations): the ambiguity is flagged for downstream processing.
  c. the peptide does not match any sequence. In this case, the mass is theoretically calculated, and the algorithm slides over all protein sequences (i.e. not only considering tryptic peptides), looking for an isobaric match. This needs to exactly match the peptide mass accompanied with combinatorial masses of the PTMs under investigation. Then, if the original peptide ends in a K or an R, it filters out only those candidates, otherwise all candidates (tryptic and semi-tryptic alike) are retained. All potential peptide-protein sequence matches are returned and aligned, allowing to consider gaps in either the measured or the theoretical peptide sequence. Then the algorithm will try to resolve each mismatch or gap in the alignment with the isobaric matrix. Only peptides that can be turned into the protein sequence match by introducing local isobaric switches are retained.
4. The list of candidate peptides is purified by discarding:
  a. All peptides that are isobaric to trypsin or LysC peptides
  b. All single hit wonders (1 peptide to 1 protein).
  c. All flagged ambiguous peptides if no evidence for them is found in another species in the database, thus removing bias towards more completely sequenced close related species.
5. Building the “collagen taxonomic” tree: To cope with the annotation bias of the proteins in the database, the algorithm next aligns (BLOSUM90) and calculates the distance between all probable proteins that are still under consideration. These distances are used to build the collagen tree.
6. Building the NCBI taxonomic tree: Next, the species taxonomic tree is built according to NCBI taxonomy, using only the peptides, and therefore species, still under consideration. The algorithm starts at the last common ancestor of all species that are under consideration. At every branching point, the algorithm looks to see if there is a difference in protein and/or peptide content between two branches. At multifurcations, bifurcation is “enforced” through the creation of a pseudo-branch that contains all the species most related and bifurcates that away from the species that is least related. Note that for this, only peptide candidates in the data are considered, i.e. not genetic relatedness. So, each iteration 2 branches are considered; when a branch is considered a subset of another branch, this branch is discarded, when both show signs of uniqueness, both are retained, and the sample is considered to be a mixture. The algorithm halts at a higher taxonomic level when no difference between the branches is found (see Figure x for what this implies to the user). After each potential endpoint has been found, the algorithm filters out all species which have less than 10% coverage of the total amount of collagen peptides in the sample. Also, all species are discarded that show to be a <80% subset of a mixture of two other species.
7. Now, for each of the possible species a Bray Curtis score is assigned based on the peptide content of the species compared against the total collagen related peptide list (figure/formula x in the main text). Additionally, all peptides that show signs of in-sample decay are given a lower weight during scoring.
8. The algorithm generates an interactive sunburst plot for easy visualization of the results, as well as a CSV result file. This sunburst plot depicts all the species that could not be filtered out during the taxonomic classification.
9. Finally, a rescoring is done. Therefore, from the result file the top hits are reconsidered and rescored by the Bray Curtis score while considering only this subset of peptides *de facto* changing the ration of the enumerator and denominator and resulting in a more resolved scoring. Such rescoring has the added benefit of helping to resolve whether the sample is a genetic or a physical mixture, based on uniqueness amongst the top scoring hits (as described in the main text). These results are visualized as a line plot of the score and the rescore as well as a barplot showing the overlap of unique peptides. Common peptides between the top results are discarded for the rescoring.

### Peptide identification

Raw datafiles were peak-picked by the MSConvert peak-picking algorithm into MGF file format (*70*). The MGF files were submitted into MASCOT Daemon (version 2.8.2, Matrix Science, London, UK), and searched against the manually curated collagen database and a Universal Contaminants database (*n*=381) (*71*) The enzyme was set to semiTrypsin with a maximum of 1 missed cleavage. Methylthio (C, +45.987721 Da) was added as a fixed modification, and Deamidation (NQ, +0.984016 Da), Oxidation (MP, +15.994915 Da) were added as variable modifications. The fragment ppm error was set to 50 ppm and the peptide error tolerance was set to 10ppm.

For species inference analysis, the Mascot search results were extracted in CSV data format restricted with significance threshold p<0.01.

### Publicly available datasets

Raw datafiles were collected from the PRIDE ProteomeXchange repository with identifiers PXD024487 (*8*), PXD031386 (*51*) and PXD042536 (*39*), and were processed in the same way as the in-house-generated DDA datasets. Morphological and analytical species classifications were downloaded from their respective papers.

## Supporting information

Database files and output overview file

ClassiCOL output files

## Acknowledgements

We would like to give special thanks to Rachel Sian Dennis for laying the groundwork for the protein extraction protocol, and giving valuable insights during the adaptation of said protocol. Also, we would like to express our gratitude to ProGenTomics (www.progentomics.be) for providing the in-house LC-MS/MS data. We thank Magnus Palmblad for providing valuable feedback on the manuscript.

## Author contributions

Conceptualization: IE, AB, SD, MD

Sample collection ADP, OUD, RMC: PR

Sample collection Scladina Cave: GA

Permissions to extract original proteomics samples: KDM, MO, VF, GA, LS, PVdP

Morphological reassessment of remains: CP, GA, KDM

Protein extraction: IE, AB

LC-MS/MS optimization: AB, SD, MD

Coding and visualization: IE, RB

Data processing: IE

Supervision: MD, DD, IDG

Optimization and validation of algorithm: IE

Writing—original draft: IE, AB, SD, MD

Writing—archeological contributions: GA, PR, IDG, CP

Writing—review & editing: all authors read and provided feedback on the manuscript

## Data and materials availability

All data are available in the main text or the supplementary materials. All original analyses were performed with ClassiCOL version 1.0.0 which is made available on Github (https://github.com/EngelsI/ClassiCOL). The in-house-generated Thermo RAW, MGF and mzTab files have been deposited in the PRIDE repository (https://www.ebi.ac.uk/pride/) with identifier PXD055222.

