## Supplementary data and figures for "ClassiCOL: LC-MS/MS analysis for ancient species Classification via Collagen peptide ambiguation"

Ian Engels et al.

**This PDF file includes:**

Supplementary figure 1: Sunburst view of Unipept output from reference Homo sapiens data. ... 3
Supplementary figure 2: Interpretation of heatmap and sunburst plot of Pecora ClassiCOL pipeline example on Rangifer tarandus sample. ... 4
Supplementary figure 3: Pongo pygmaeus identification of Rüther et al. 2022 reference dataset. 5
Supplementary figure 4: A reference dataset of Equus asinus from Rüther et al 2022 was classified as Equus quagga by ClassiCOL. ... 6
Supplementary figure 5: Parental mixture output of reference dataset of mule from Rüther et al. 2022 ... 7
Supplementary figure 6: The use of non-common collagen proteins and re-use of peptide on different taxonomic levels results in more precise classifications. ... 8
Supplementary figure 7: A proteomically determined Bos sample is more likely to be closer related to the genus Bison. ... 9
Supplementary figure 8: Analysis of the public dataset of Gilbert et al., 2024 revealed that the species of origin is more closely related to a species not included in the original papers' database. ... 10
Supplementary figure 9: COL3A1 commonly found in glue identified on ivory figurine. ... 11
Supplementary figure 10: Morphologically estimated Cervus species ADP006 turns out to be a wild boar. ... 12
Supplementary figure 11: Morphologically estimated Vulpes species ADP0014 turns out to be an otter. ... 13
Supplementary figure 12: Morphologically estimated Capreolus capreolus ADP0011 was classified as Cervus. ... 14
Supplementary figure 13: Morphologically estimated Cervus species OD024 turns out to be a goat ... 15
Supplementary figure 14: Morphologically estimated Caprinae species OD08 turns out to be a kind of deer. ... 16
Supplementary figure 15: Morphologically estimated cat OD042 turns out to be an otter. ... 17
Supplementary figure 16: Morphologically estimated Bovine species OD029 turns out to have originated from Cervinae. ... 18
Supplementary figure 17: Morphologically estimated Bovine species OD031 turns out to have originated from Cervinae, with low peptide count. ... 19
Supplementary figure 18: Fish remains OD0001, morphologically estimated as Tinca tinca is classified to the closest extant relative in the database. ... 20

Supplementary data 1: Fragment of Collagen Database overview file. 29
Supplementary data 2: Sunburst plot collection, Mascot csv files and ClassiCOL output files. (separate file) 30
Supplementary data 3: Classicol output overview table. 31
Supplementary data 4: Detailed step-by-step protein extraction protocol, 32

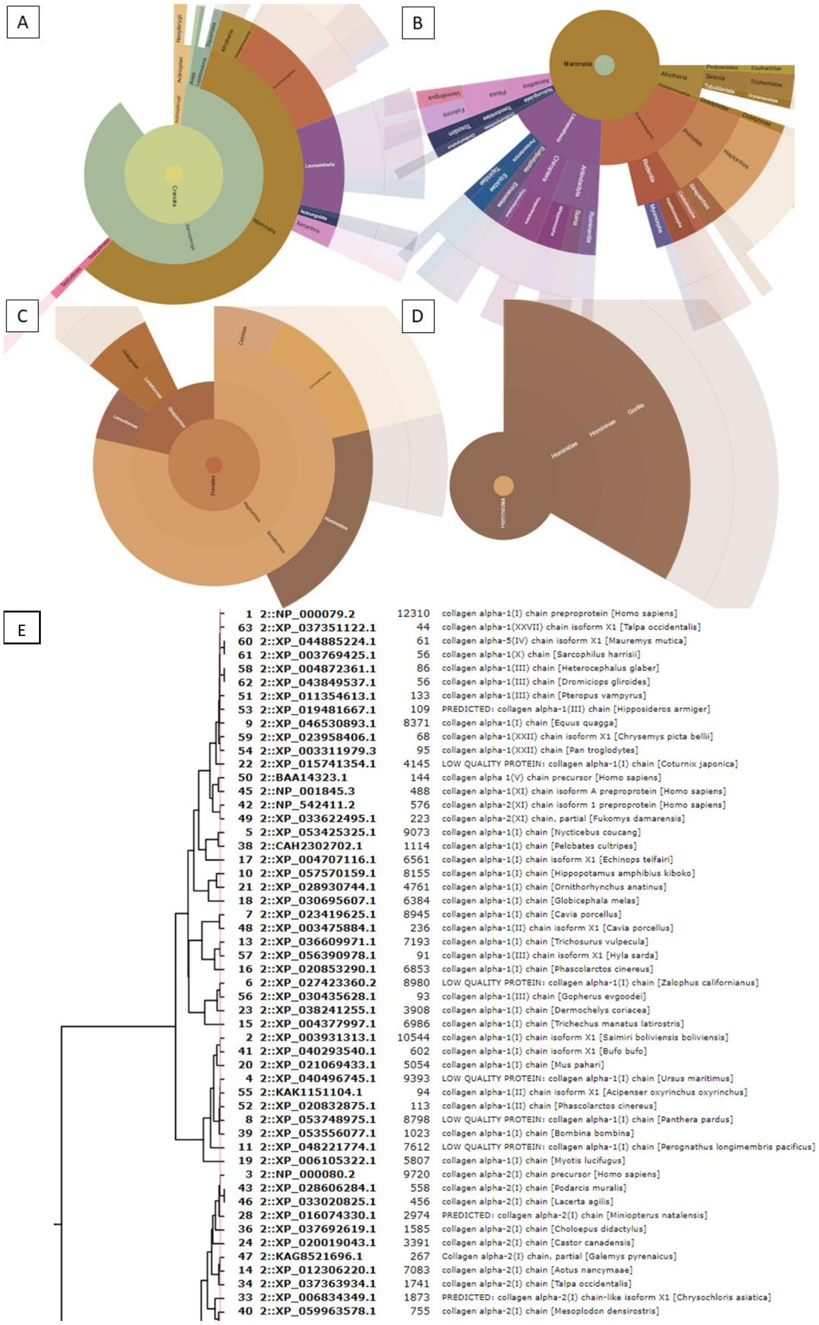

**Supplementary figure 1: Sunburst view of Unipept output from reference Homo sapiens data.** Sunburst plot was made using Unipept desktop. It visualizes the ambiguity returned by the MASCOT search engine when a database with highly similar sequences is used. From A-B the figure zooms in onto A. Craniata, B. Mammalia, C. Primates and D. Hominidae. As seen in D, Homo sapiens is not a possible outcome, although human bone was analyzed. E is an exemplary MASCOT output file of the Human reference data. When an DDA run is performed on a tryptically digested bone sample, it can be searched using a probabilistic scoring algorithm like Mascot. Still, because of small isobaric alterations between homologues collagen peptides over dozens of species, outputting the highest scoring PSM for each spectrum leads to a long list of potential organisms, effectively impairing easy annotation.

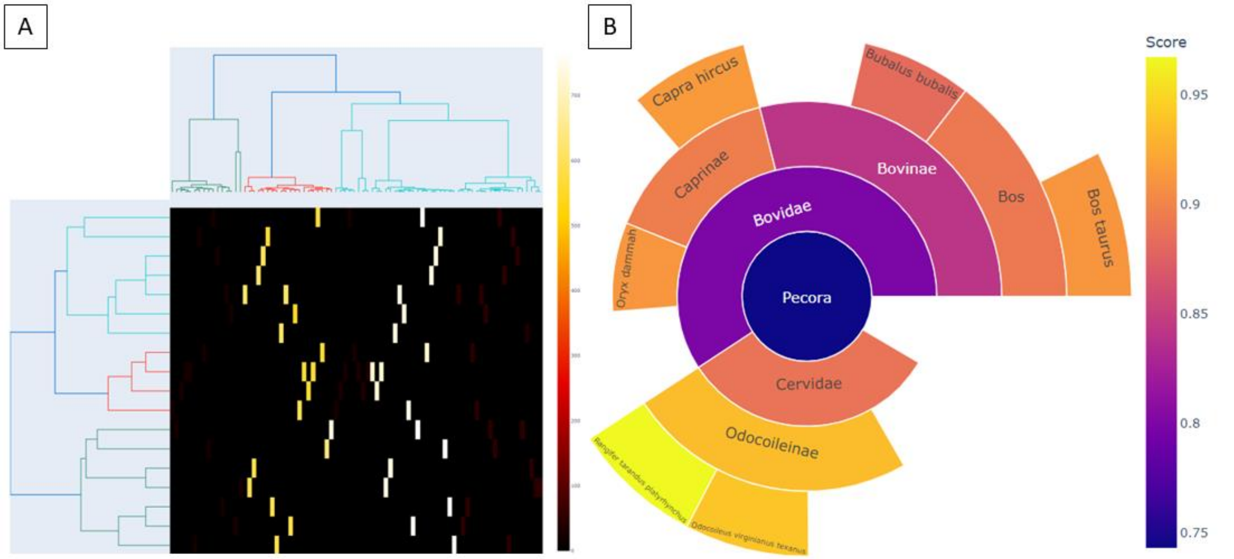

**Supplementary figure 2: Interpretation of heatmap and sunburst plot of Pecora ClassiCOL pipeline example on Rangifer tarandus sample.**

Simplified ClassiCOL output files used for ClassiCOL pipeline overview figure 2 in the main text. A. is a heatmap view with on the Y-axis the taxonomic tree of Pecora species in the database, on the X-axis a homology based tree of the different collagen sequences represented by at least 2 peptides. The heatmap is colored by number of unique peptides per protein. B. The sunburst output plot of the Pecora family. Here the different taxa are colored by score assigned by ClassiCOL. Both figures are interactive and can be found in the Supplementary data folder 'Additional_figures'.

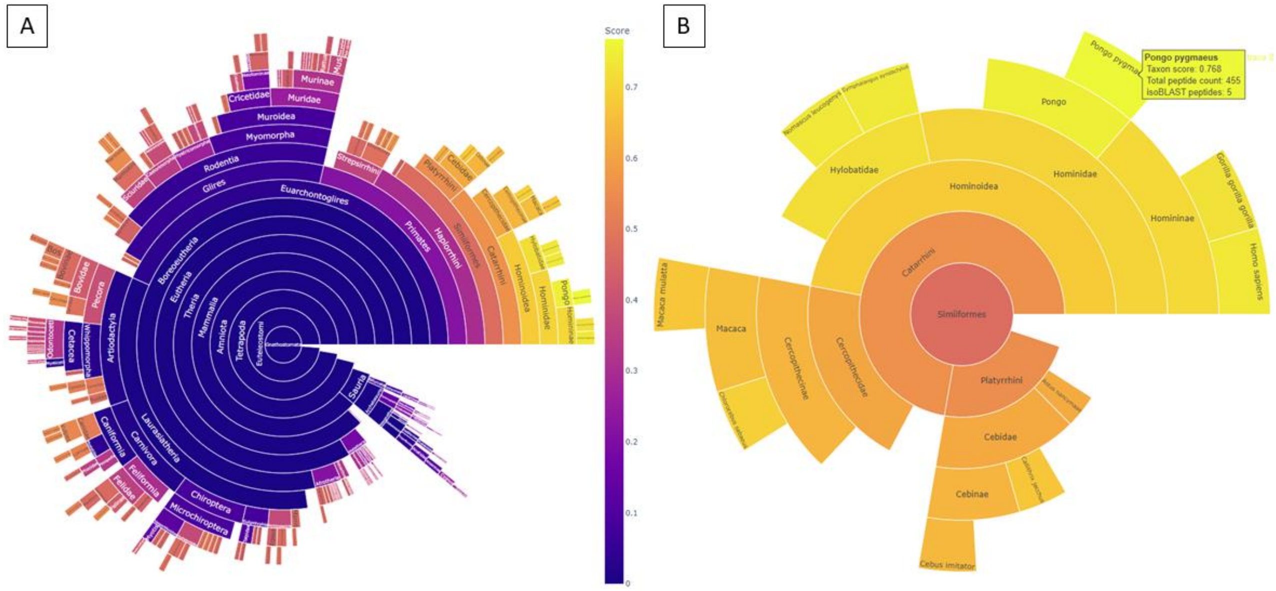

**Supplementary figure 3: Pongo pygmaeus identification of Rüther et al. 2022 reference dataset.**
Here the ClassiCOL output of file 20200710_EXPL2_Evo2_PR_SA_100spd_DDA_Ancient_PrimateBone_Pongo-pygmaeus_Orang1 is shown. This demonstrates that the selection of species to be included in the database influences the final taxonomic output. In the original paper the proteomics classification resulted in a different species in the genus Pongo due to lack of protein sequences of the morphological identification. A. Sunburst plot showing the general overview of the ClassiCOL output. B. The same sunburst plot as in A. zoomed in to Simiformes, top resulting identification shown by the yellow textbox.

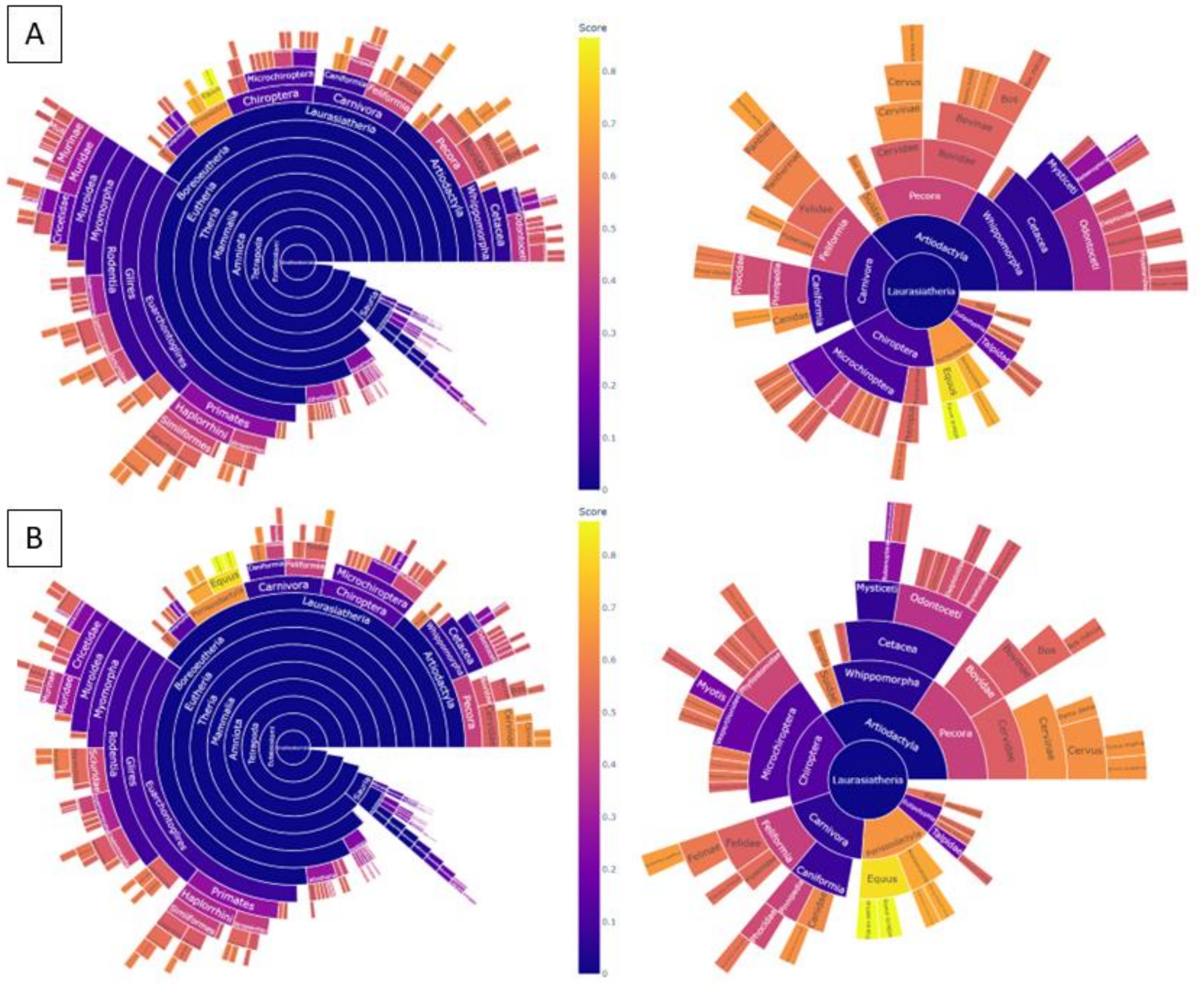

**Supplementary figure 4: A reference dataset of Equus asinus from Rüther et al 2022 was classified as Equus quagga by ClassiCOL.**

A. ClassiCOL output of reference dataset 20200710_EXPL2_Evo2_PR_SA_100spd_DDA_Ancient_KnownBone_Equus-asinus_P23. Left = general overview figure of the Classicol output. Right=Zoomed in on Laurasiatheria. B. ClassiCOL output of reference dataset 20200710_EXPL2_Evo2_PR_SA_100spd_DDA_Ancient_KnownBone_Equus-asinus_P22. Assignation of Equus asinus rather than Equus quagga in figure A. Is due to a peptide shared between Equus quagga and Equus caballus.In figure B. no distinctive peptides were found that can differentiate between Equus asinus and Equus quagga.

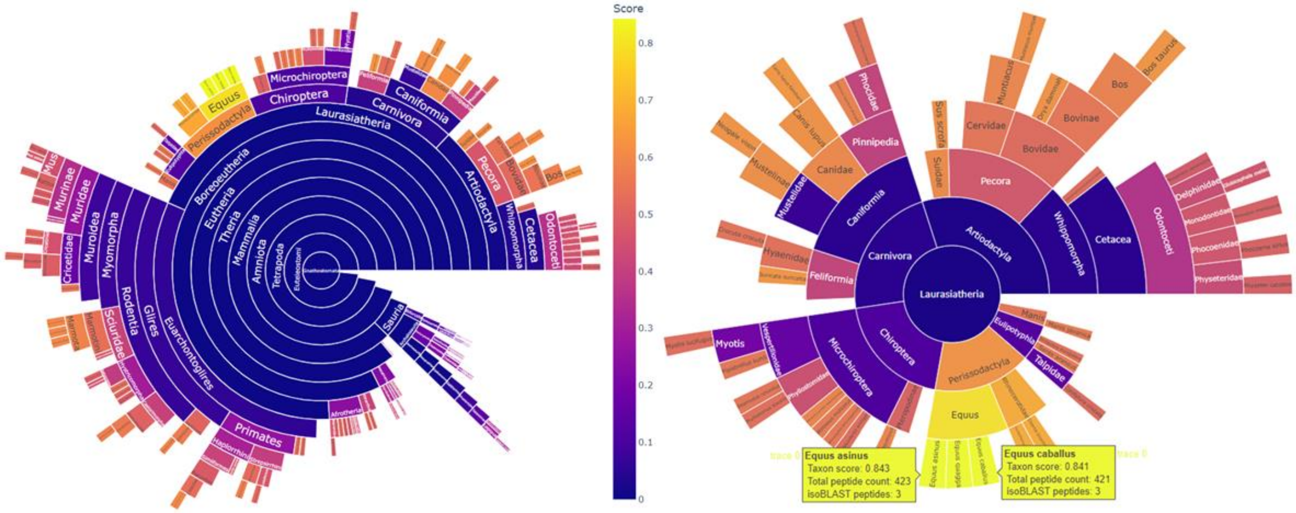

**Supplementary figure 5: Parental mixture output of reference dataset of mule from Rüther et al. 2022.**
The left figure shows the general overview output of ClassiCOL for the reference dataset file 20200710_EXPL2_Evo2_PR_SA_100spd_DDA_Ancient_KnownBone_Equus-Asinus-X-Caballus_P35. The right figure is zoomed in on the Laurasiatheria, with Equus caballus and Equus asinus highlighted. This shows that Equus caballus is scored differently to Equus asinus, but both were retained as possible outcomes. This is due to the finding of uniqueness in peptides for both species. This is in accordance with the specimen being derived from both species of horse i.e. a parental mixture.

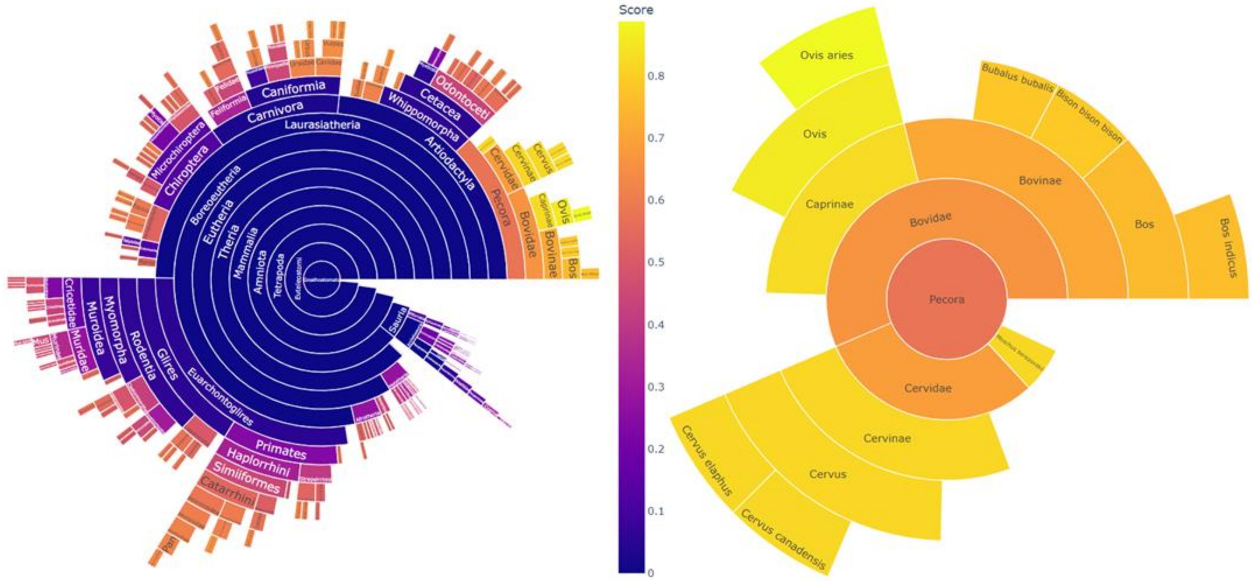

**Supplementary figure 6: The use of non-common collagen proteins and re-use of peptide on different taxonomic levels results in more precise classifications.**

Here a sunburst output is shown of Salpetermosen samples from Rüther et al 2022, file 20200709_EXPL2_Evo2_IH_SA_100spd_DDA_Ancient_unknown_Salpetermosen_Bone-8326. The original paper classified the sample at the Caprinae level, after ClassiCOL analysis Ovis aries was returned as most likely species. Carpra hircus was not in the possibility list. This was in accordance with the original paper's listed morphology for this specific sample.

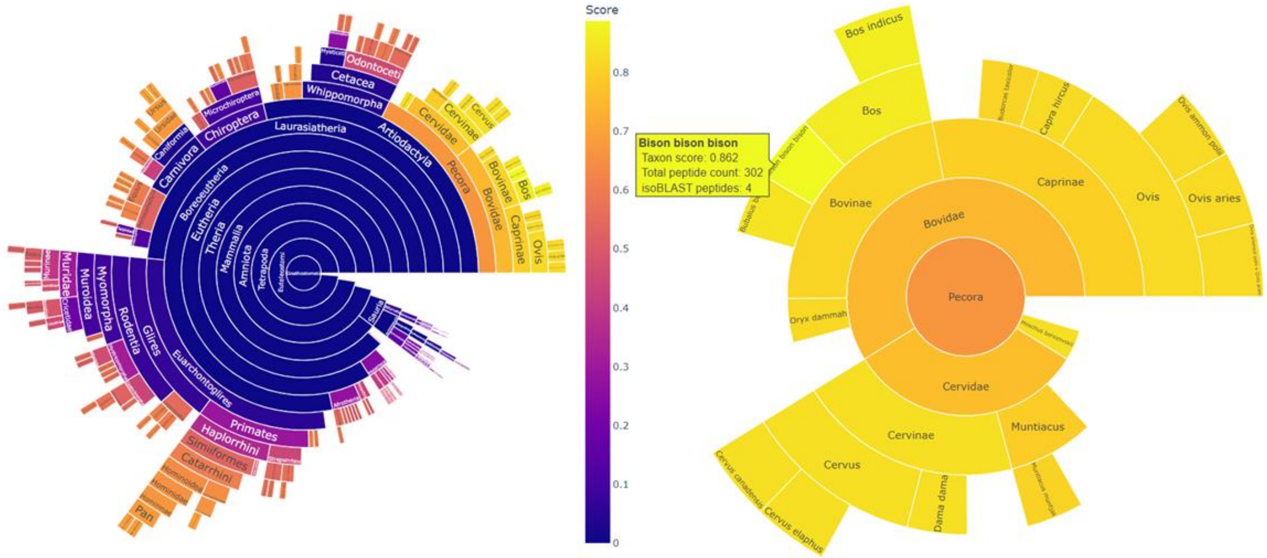

**Supplementary figure 7: A proteomically determined Bos sample is more likely to be closer related to the genus Bison.**

Here a sunburst output is shown of Salpetermosen samples from Rüther et al 2022, file 20200709_EXPL2_Evo2_IH_SA_100spd_DDA_Ancient_unknown_Salpetermosen_Bone-5306. Unlike the proteomics identification of the original paper ClassiCOL classified the specimen as having originated from Bison rather than a species of Bos.

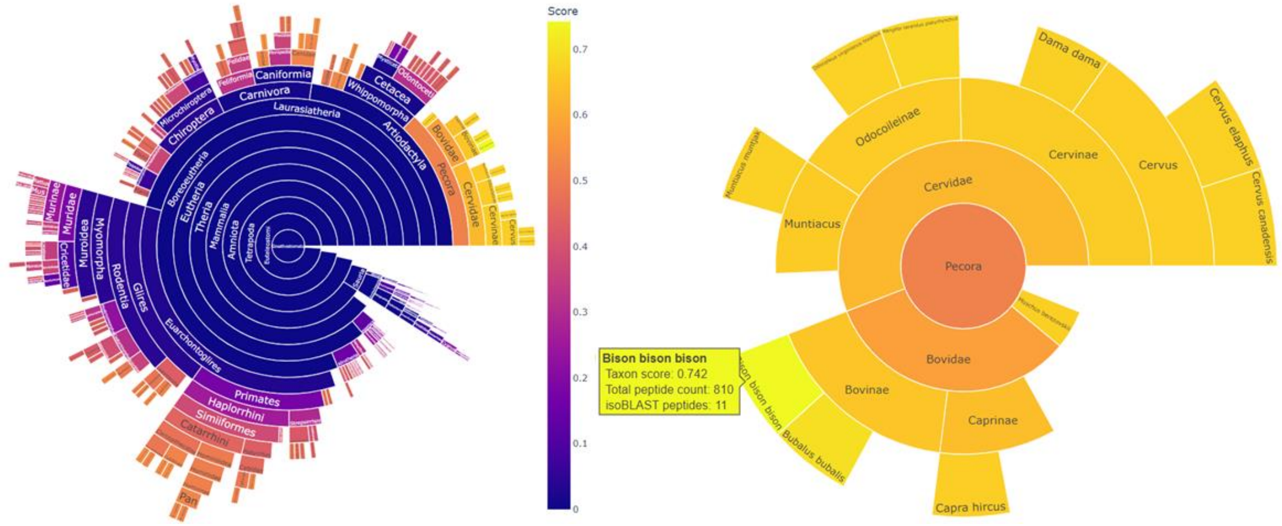

**Supplementary figure 8: Analysis of the public dataset of Gilbert et al., 2024 revealed that the species of origin is more closely related to a species not included in the original papers' database.**
Here a sunburst plot is show from the ClassiCOL output on a publicly available file from Gilbert et al.2024 file 15-PUSHH220808_Virginal_Akey_11-176-1. In the original paper Bison was not considered in the database. By using the extensive ClassiCOL collagen database Bison bison is the most closely related species in the database.

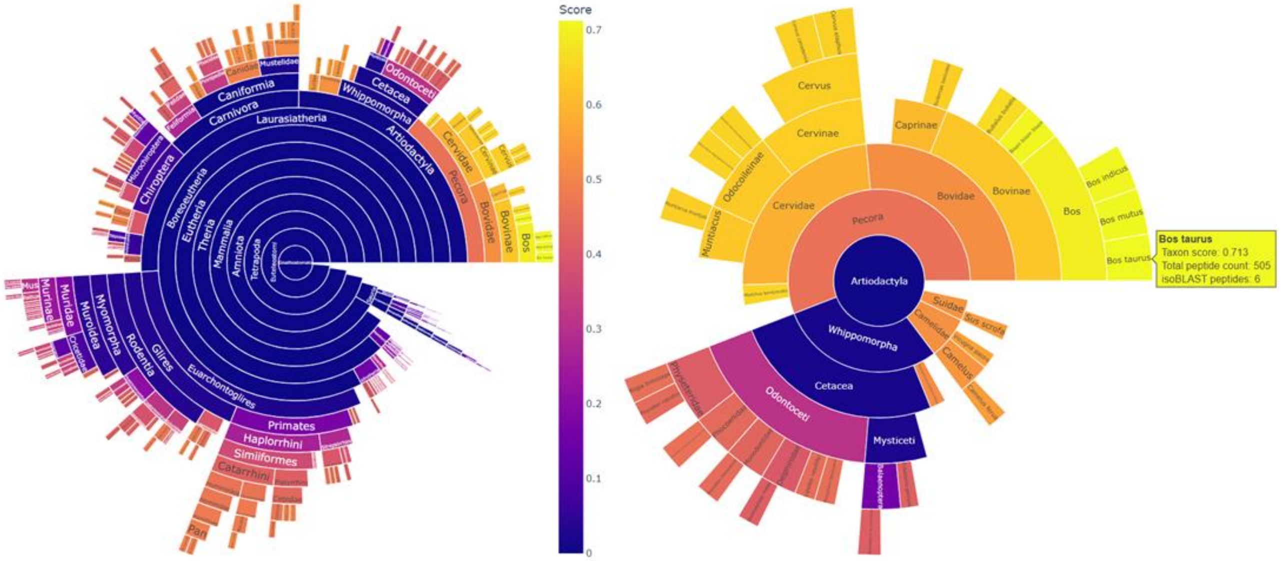

**Supplementary figure 9: COL3A1 commonly found in glue identified on ivory figurine.**
Here a sunburst plot is show from the ClassiCOL output on a publicly available file from Gilbert et al.2024 file 4-PUSHH220916_66-99-50_Figurine. As stated in the original paper, the sample collected most likely contained animal glue traces. In particular, glue derived from Bos taurus origin, in agreement with the original proteomics identification.

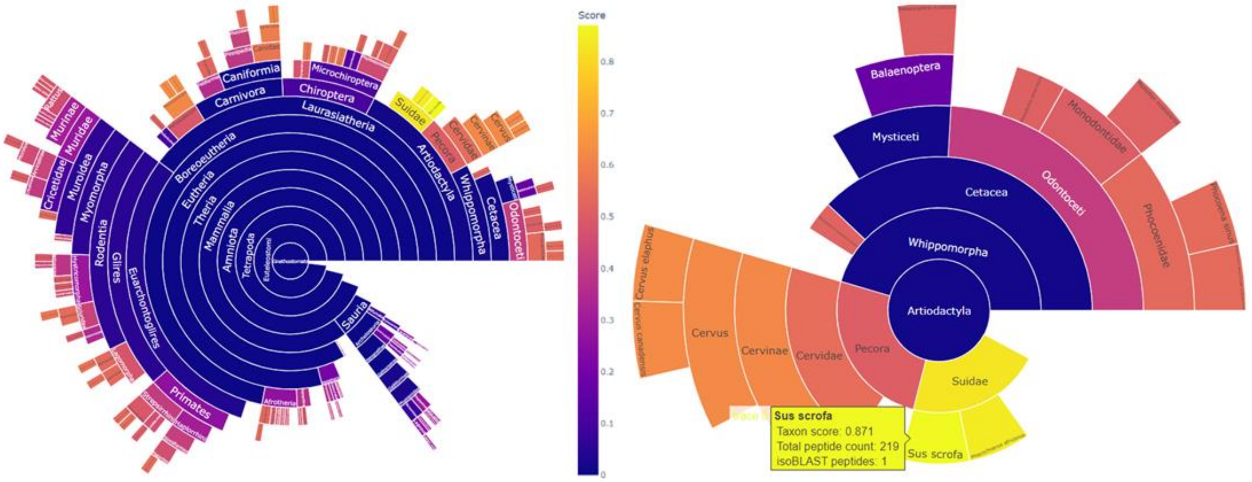

**Supplementary figure 10: Morphologically estimated Cervus species ADP006 turns out to be a wild boar.** In-house processed sample ADP006 was originally estimated to have derived from a kind of cervid. The ClassiCOL analysis, here represented by sunburst plot in its entirety left and zoomed in at taxonomic level containing morphology and algorithmic classification on the right, classified the sample as Sus scrofa. The Sus scrofa identification was confirmed via morphological re-estimation.

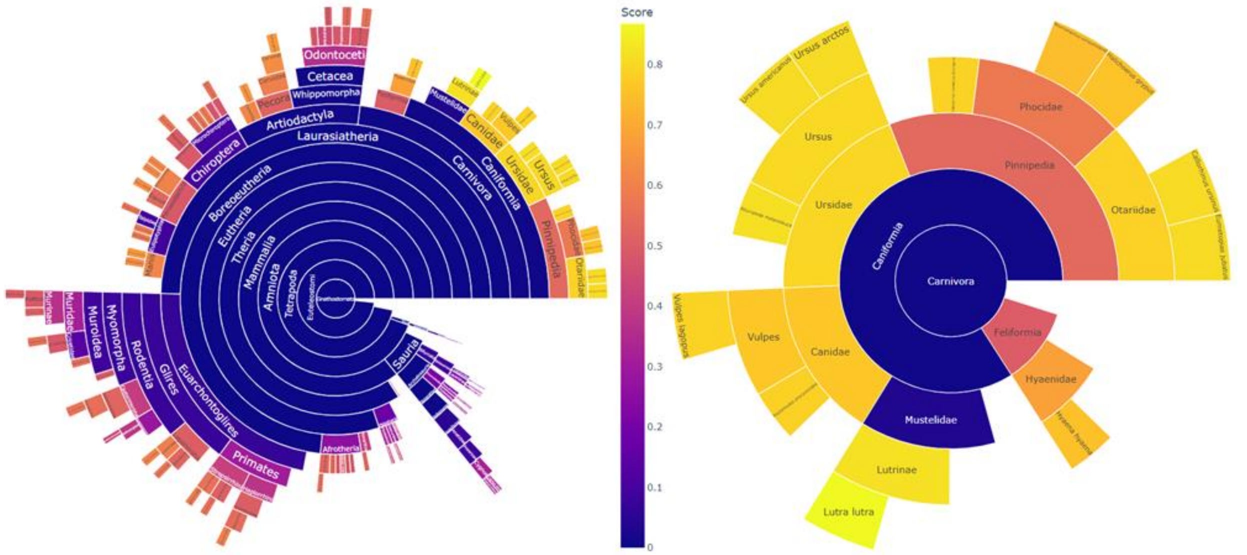

**Supplementary figure 11: Morphologically estimated Vulpes species ADP0014 turns out to be an otter.** In-house processed sample ADP0014 was originally estimated to have derived from Vulpes vulpes. The ClassiCOL analysis, here represented by sunburst plot in its entirety left and zoomed in at taxonomic level containing morphology and algorithmic classification on the right, classified the sample as Lutra lutra. The Lutra lutra identification was confirmed via morphological re-estimation.

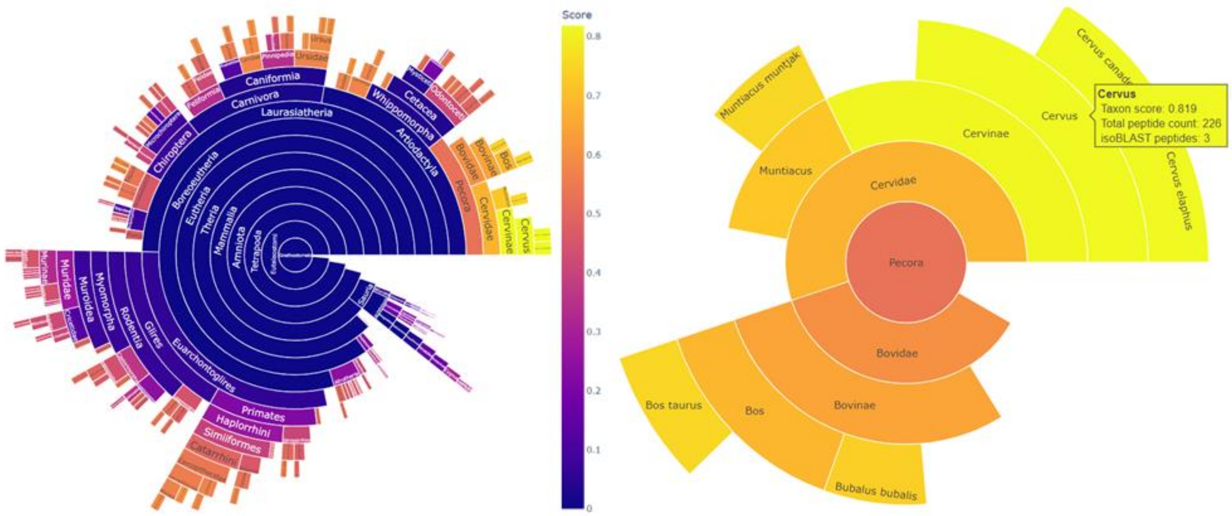

**Supplementary figure 12: Morphologically estimated Capreolus capreolus ADP0011 was classified as Cervus.** In-house processed sample ADP0011 was originally estimated to have derived from Capreolus capreolus. The ClassiCOL analysis, here represented by sunburst plot in its entirety (left) and zoomed in at taxonomic level containing morphology and algorithmic classification (right), classified the sample as Cervus. The Cervus identification was deemed improbable due to morphological traits that differ between the two genera.

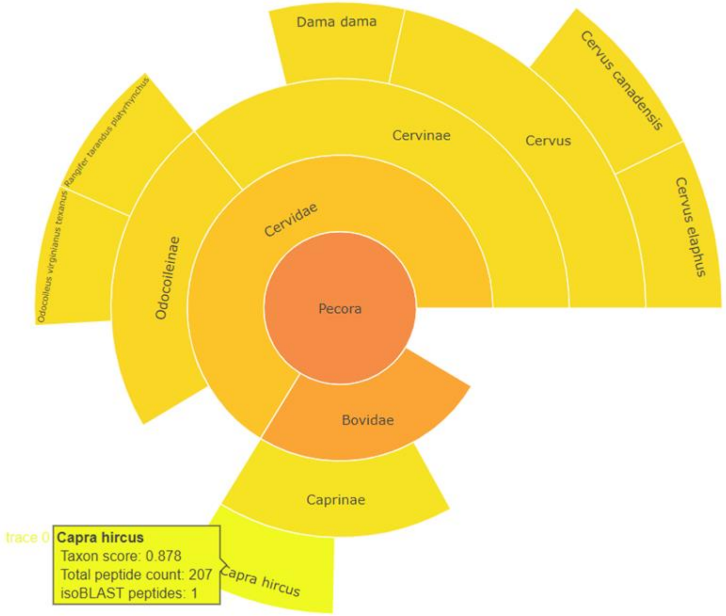

**Supplementary figure 13: Morphologically estimated Cervus species OD024 turns out to be a goat**
In-house processed sample OD024 was originally estimated to have derived from the genus Cervus. The ClassiCOL analysis, here represented by sunburst plot zoomed in at taxonomic level containing morphology and algorithmic classification, classified the sample as Capra hircus. The Capra hircus identification could not be morphologically confirmed due to the state of the bone fragment.

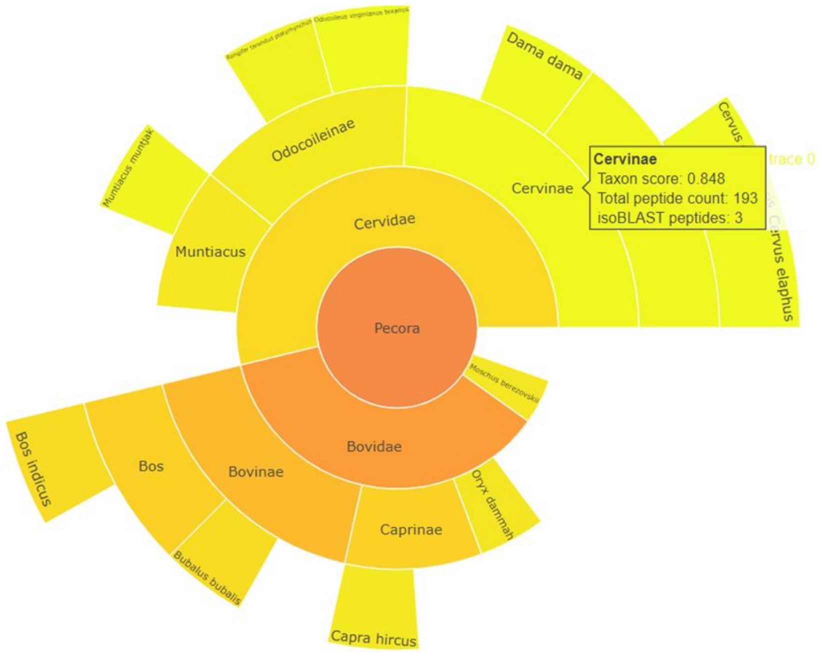

**Supplementary figure 14: Morphologically estimated Caprinae species OD08 turns out to be a kind of deer.** In-house processed sample OD08 was originally estimated to have derived from Caprinae. The ClassiCOL analysis, here represented by sunburst plot zoomed in at taxonomic level containing morphology and algorithmic classification, classified the sample as Cervinae. This identification could not be morphologically confirmed due to the state of the bone fragment.

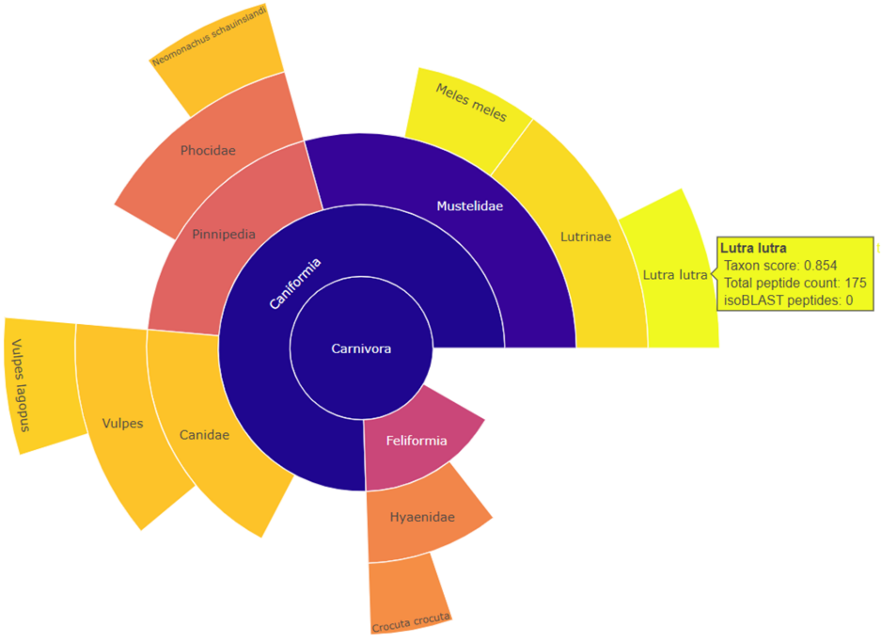

**Supplementary figure 15: Morphologically estimated cat OD042 turns out to be an otter.**
In-house processed sample OD042 was originally estimated to have derived from Felis catus. The ClassiCOL analysis, here represented by sunburst plot zoomed in at taxonomic level containing morphology and algorithmic classification, classified the sample as Lutra lutra. The morphologically estimated species was not found in the list of possible outcomes. The Lutra lutra identification could not be morphologically confirmed due to the state of the bone fragment.

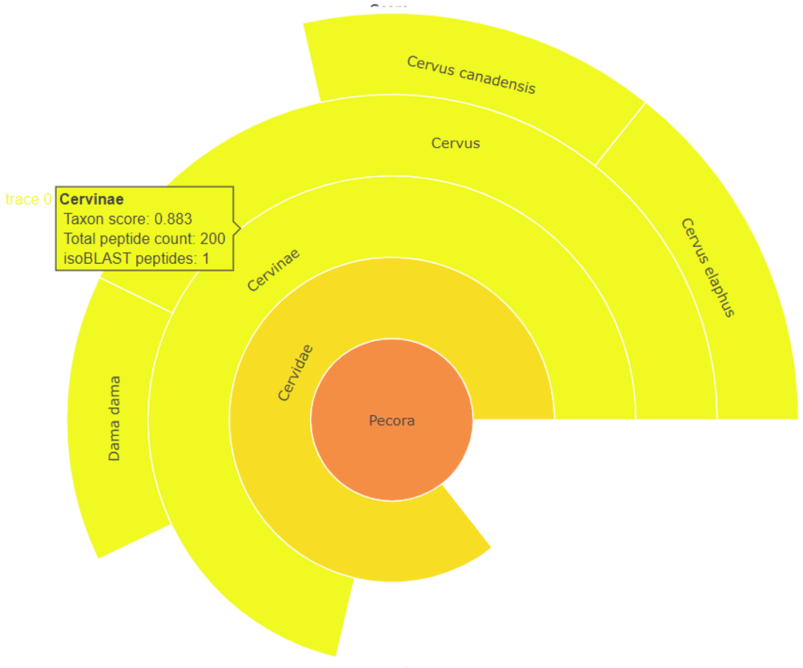

**Supplementary figure 16: Morphologically estimated Bovine species OD029 turns out to have originated from Cervinae.**

In-house processed sample OD029 was originally estimated to have derived from Bovinae. The ClassiCOL analysis, here represented by sunburst plot zoomed in at taxonomic level containing morphology and algorithmic classification, classified the sample as Cervinae. The morphologically estimated species was not found in the list of possible outcomes. This identification could not be morphologically confirmed due to the state of the bone fragment.

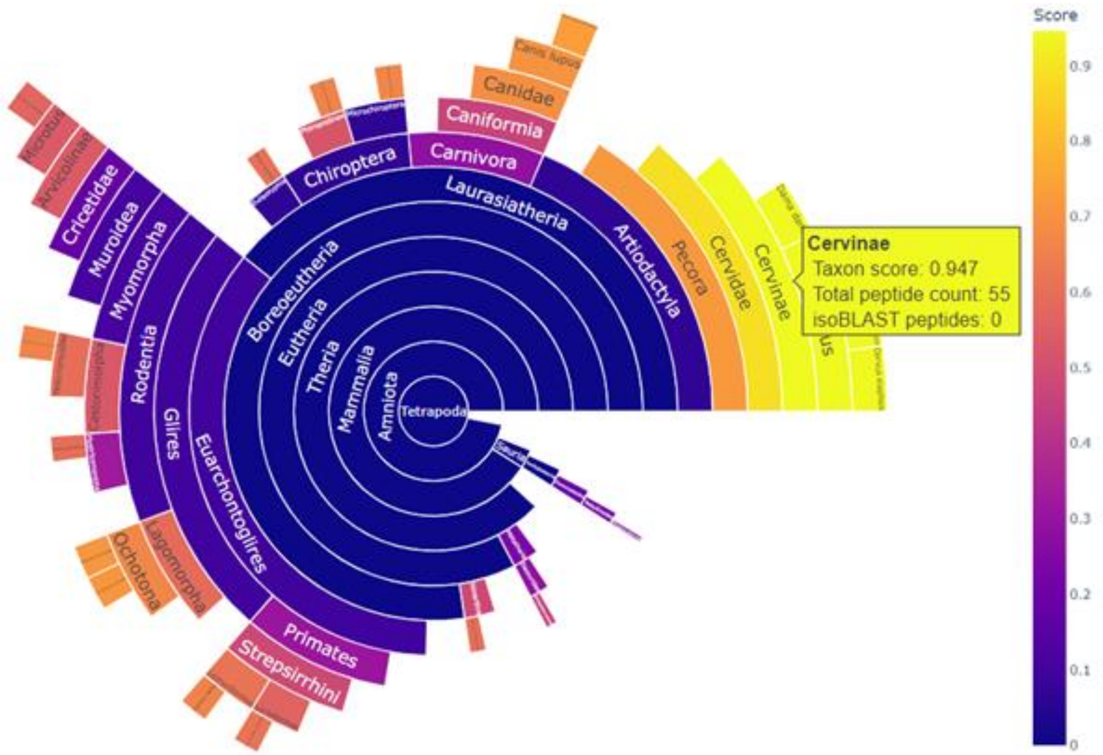

**Supplementary figure 17: Morphologically estimated Bovine species OD031 turns out to have originated from Cervinae, with low peptide count.**
In-house processed sample OD031 was originally estimated to have derived from the genus Bos. The ClassiCOL analysis, here represented by sunburst plot zoomed in at taxonomic level containing morphology and algorithmic classification, classified the sample as Cervinae. The morphologically estimated species was not found in the list of possible outcomes. This identification could not be morphologically confirmed due to the state of the bone fragment. Additionally, the number of peptides found in the sample covered the matching protein sequence to a low percentage. This could be the reason for higher taxonomic classification**.**

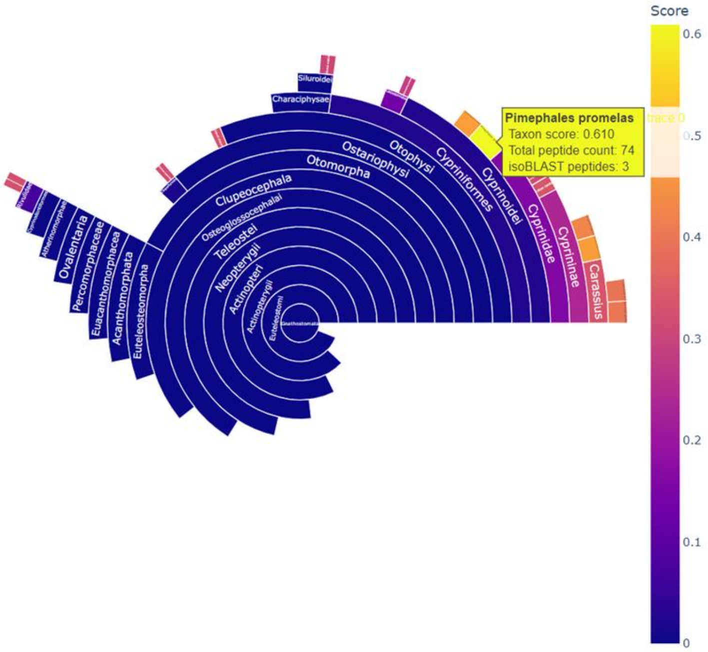

**Supplementary figure 18: Fish remains OD0001, morphologically estimated as Tinca tinca is classified to the closest extant relative in the database.**

In-house processed sample OD0001 was originally estimated to have derived from Tinca tinca. The ClassiCOL analysis, here represented by sunburst plot, classified the sample as to its closest extant relative in the collagen database. This identification could not be morphologically confirmed due to the state of the bone fragment.

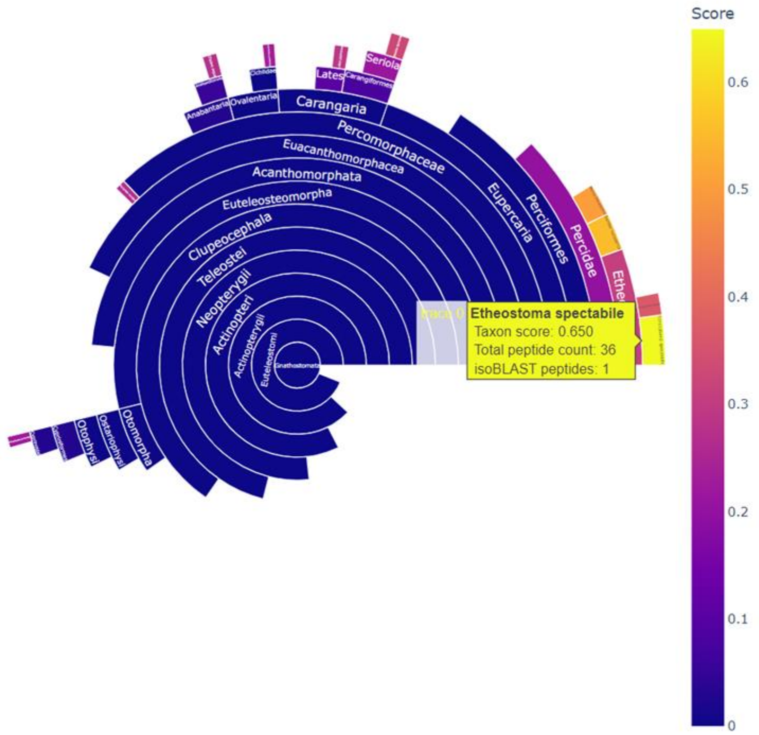

**Supplementary figure 19: Fish remains OD0002, morphologically estimated as Perca fluviatus is classified to the closest extant relative in the database.**

In-house processed sample OD0001 was originally estimated to have derived from Perca fluviatus. The ClassiCOL analysis, here represented by sunburst plot, classified the sample to another species in the family Percidae. This could suggest a genetic mixture of extant species of Percidae in the database. In other words, the true species is not present in the database.

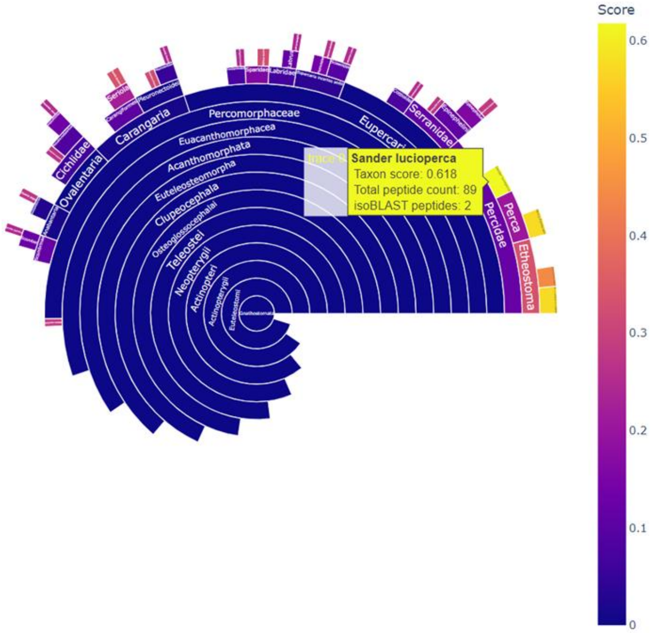

**Supplementary figure 20: Fish remains OD0003, morphologically estimated as a kind of fish is classified to have originated from the family Percidae.**

In-house processed sample OD0003 was originally estimated to have derived from a kind of fish. The ClassiCOL analysis, here represented by sunburst plot, classified the sample to Sander lucioperca. The results also suggest a genetic mixture of extant species of Percidae in the database. In other words, the true species could potentially not be present in the database. This identification could not be morphologically confirmed due to the state of the bone fragment.

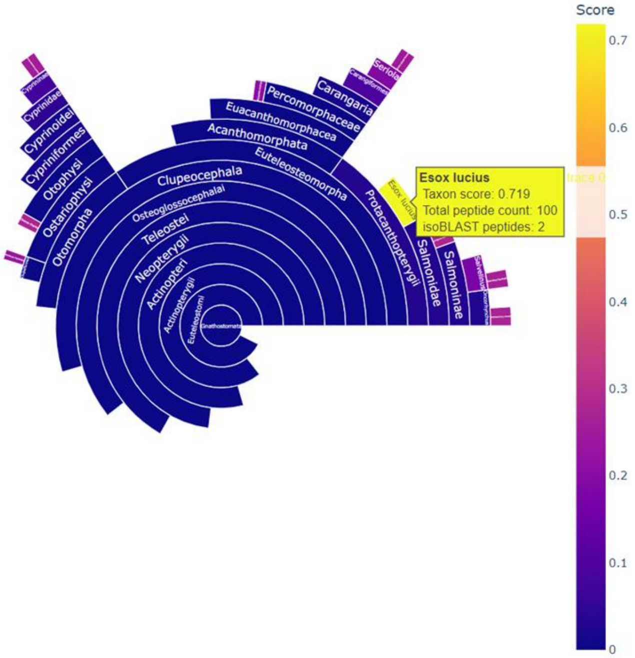

**Supplementary figure 21: Fish remain OD0007, morphologically estimated as Esox lucius is classified to the correct species.**
In-house processed sample OD0007 was originally estimated to have derived from Esox lucius. The ClassiCOL analysis, here represented by sunburst plot, classified the sample also as Esox Lucius, with a high score.

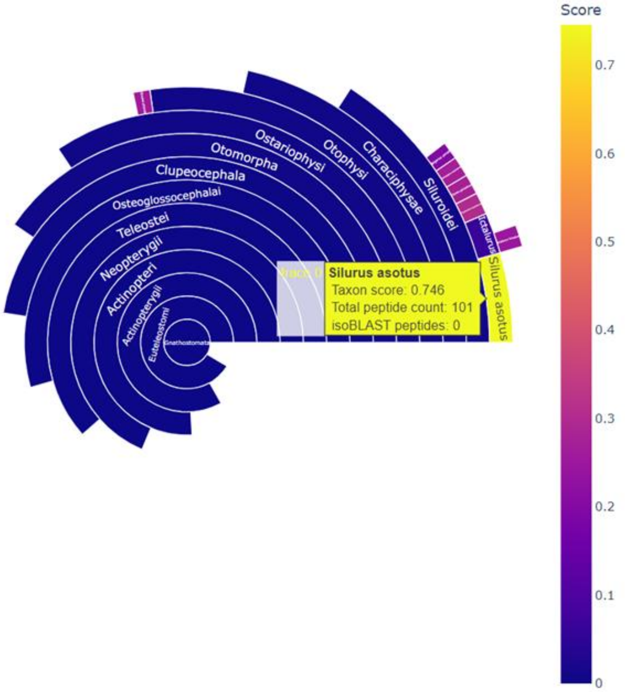

**Supplementary figure 22: Fish remains OD0004, morphologically estimated as Silurus glanis is classified to another member of the genus in the database.**
In-house processed sample OD0004 was originally estimated to have derived from Silurus glanis. The ClassiCOL analysis, here represented by sunburst plot, classified the sample as Silurus asotus. This species, also from the genus Silurus, is the closest extant relative in the database.

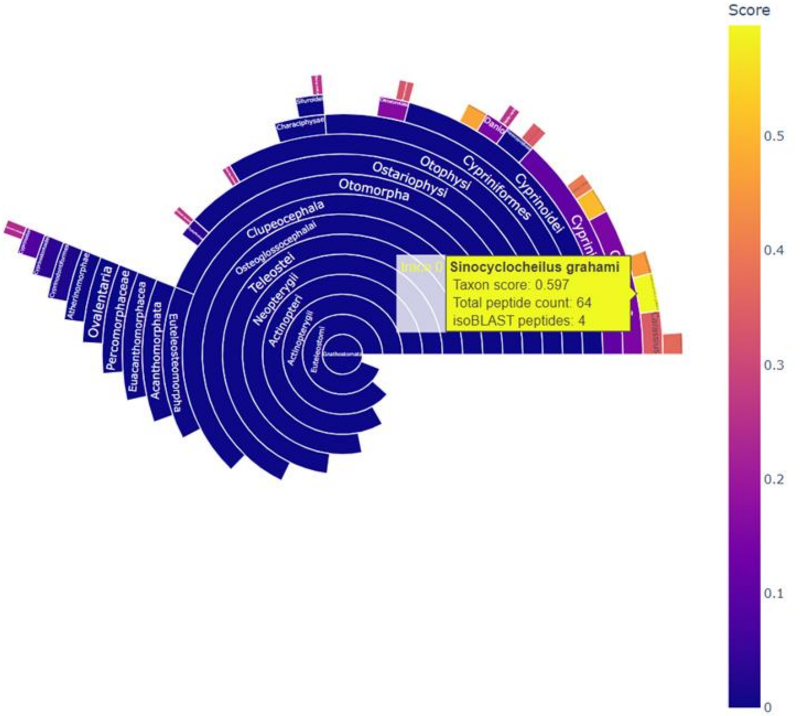

**Supplementary figure 23: Fish remains OD0009, morphologically estimated as Barbus barbus is classified to the closest extant relative in the database.**
In-house processed sample OD0009 was originally estimated to have derived from Barbus barbus. The ClassiCOL analysis, here represented by sunburst plot, classified the sample as to another species in the family Cyprinoidae, with Sinocyclocheillus graham as the highest match. This could suggest a genetic mixture of extant species of Cyprinoidae in the database, with the top scoring hit being the closest related to the true species.

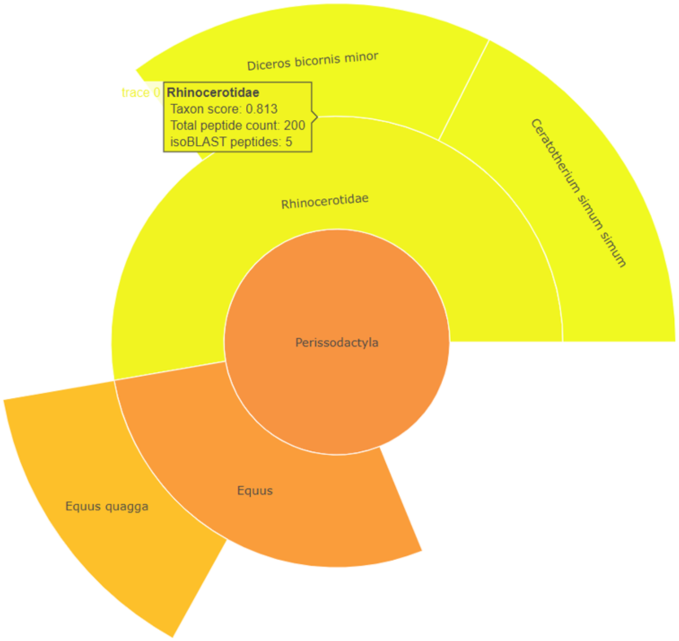

**Supplementary figure 24: Woolly rhino (SC1995-279-475) matches to closest extant rhinoceros genus in the database.**
In-house processed sample SC1995-279-475 was originally estimated to have derived from Coelodonta antiquus. The ClassiCOL analysis, here represented by sunburst plot zoomed in at taxonomic level containing morphology and algorithmic classification, classified the sample to Rhinocerotidae. This genus is the closest extant genus in the database, which contains 2 extant species. However no uniqueness to either of the two species was found.

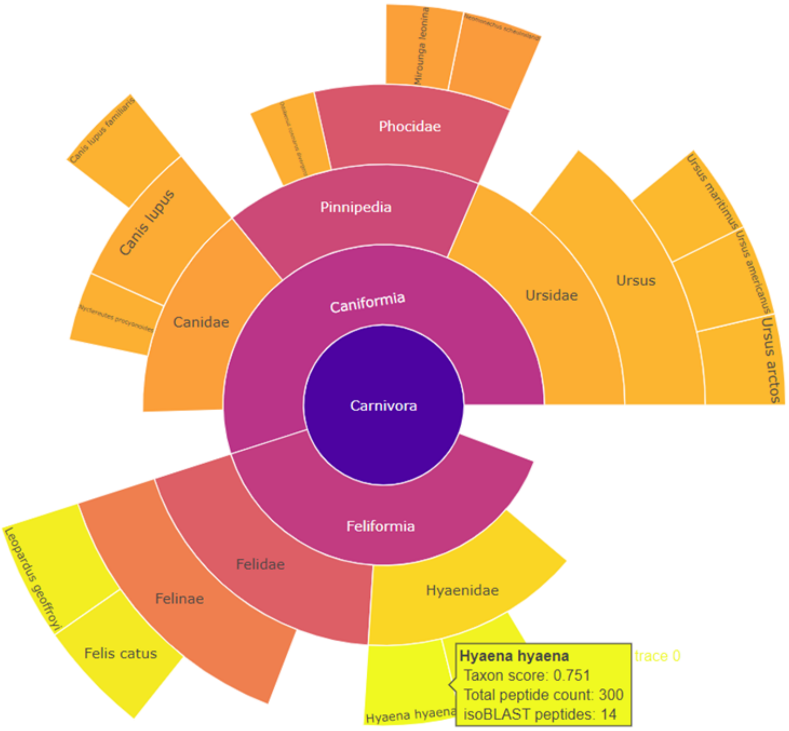

**Supplementary figure 25: Morphologically estimated lynx (SC2002-699-5) turn out to be a hyena cub.** In-house processed sample SC2002-699-5 was originally estimated to have derived from Lynx lynx. The ClassiCOL analysis, here represented by sunburst plot zoomed in at taxonomic level containing morphology and algorithmic classification, classified the sample as Hyaena hyaena. Morphological re-estimation of the bone confirmed this identification.

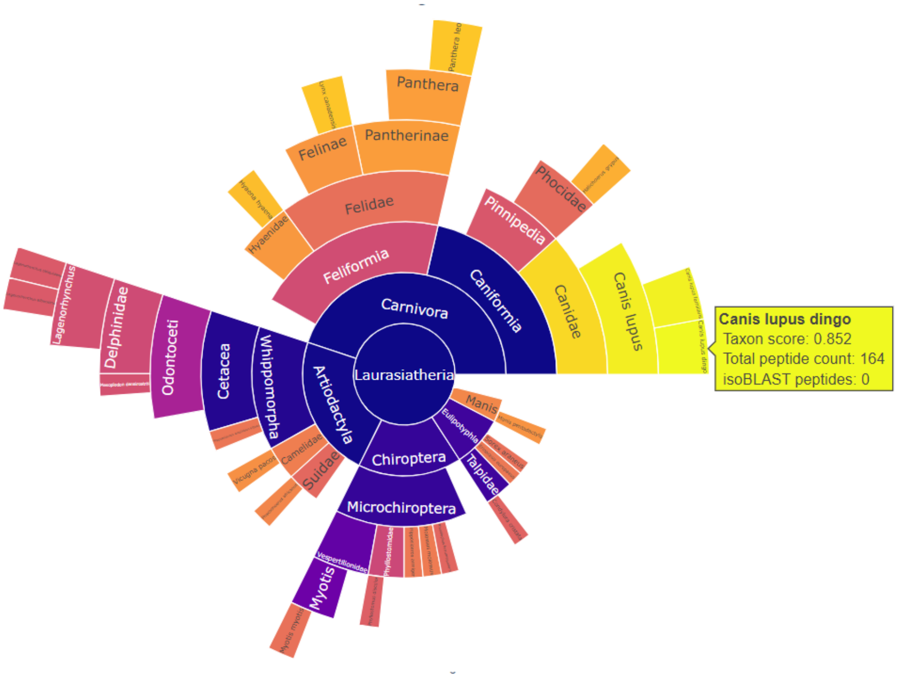

**Supplementary figure 26: Mophologically estimated Rupicapra rupicapra (SC1985-543-3) turns out to be a juvenile wolf.**

In-house processed sample SC1985-543-3 was originally estimated to have derived from Rupicapra rupicapra. The ClassiCOL analysis, here represented by sunburst plot zoomed in at taxonomic level containing morphology and algorithmic classification, classified the sample as Canis lupus dingo. Morphological re-estimation of the bone confirmed this identification. Additionally, the results show signs of genetic mixture suggesting that the samples originated from Canis lupus.

| animal | common name | Collagen A1 (I) | collagen alpha 1 (II) | collagen alpha 1 (III) | collagen alpha 1 (IV) |
|---|---|---|---|---|---|
| Acipenser ruthenus | sterlet sturgeon | X | X | X | X |
| Acipenser oxyrinchus oxyrinchus | Atlantic sturgeon | X | X | | X |
| Anas platyrhynchos | mallard duck | X | X | X | X |
| Anguilla anguilla | european eel | X | X | X | X |
| Anser cygnoides | swan goose | X | X | X | X |
| Arvicola amphibius | european water vole | X | X | X | X |
| Bison bison bison | Bison | X | X | X | X |
| Bos taurus | domestic cow | X | X | X | X |
| Bubalus bubalis | domestic water buffalo | X | X | X | X |
| Canis lupus familiaris | domestic dog | X | X | X | X |
| Capra hircus | goat | X | X | X | X |
| Castor canadensis | american beaver | X | X | X | X |
| Cervus canadensis | elk | X | X | X | X |
| Cervus elaphus | red deer | X | X | X | X |
| Columba livia | rock dove | X | X | X | X |
| Coturnix japonica | japanese quail | X | X | X | X |
| Cygnus olor | mute swan | X | X | X | X |
| Cyprinus carpio | carp fish | X | X | | X |
| Dicentrarchus labrax | european seabass | X | X | X | X |

**Supplementary data 1: Fragment of Collagen Database overview file.**
The Bone_db.xlsx document contains the scientific names alongside the common name of all species included in the ClassiCOL analysis up to August 2024. All protein sequences that were found in either NCBI or Uniprot are shown in green. Proteins for which no sequences were found are shown in red.

**Supplementary data 2: Sunburst plot collection, Mascot csv files and ClassiCOL output files. (separate file)**

All data (in-house processed and public data) analyzed via ClassiCOL return 2 main files (i) an interactive sunburst plot, which give a visual representation of all collagen related peptides in the sample and in which species these can be found. All the sunburst plots can be found in the folder 'ZooMS_output_files > SunburstPlots' in this folder all public data outputs are sorted by their paper and our in-house processed samples can be found under 'Mesolithic_samples' containing the outputs from ADP, RMC and OUD. The folder 'Scladina' contains our in-house processed data from the Scladina Cave site. (ii) Similarly, the folder 'ZooMS_output_files > ZooMS_result_files' contain each output csv with all possible species ranked to score with their corresponding proteins, peptides unique -and isobaric peptides. Similarly, all Mascot *.csv output files are available in the folder mascot_result_files in the folders for each of the archeological sites.

| raw file name Extinct collection (Bray et al. 2022) | Morphology (according to the paper) | Proteomics result paper | ClassiCOL output | Total amount of peptides | ClassiCOL score |
|---|---|---|---|---|---|
| sc-1_O | Panthera spelaea | Felidae | Panthera (Panthera leo/tigris/unc | 676 | 0.75177962 |
| sc-1-D | Panthera spelaea | Felidae | Panthera (Panthera leo/tigris) | 965 | 0.6600242 |
| sc-2_O | Panthera spelaea | Felidae | Panthera (Panthera leo/tigris/unc | 799 | 0.73358087 |
| sc-2_D | Panthera spelaea | Felidae | Panthera (Panthera leo/tigris/unc | 885 | 0.73333561 |
| sc-3_O | Panthera spelaea | Felidae | Panthera (Panthera leo/tigris/unc | 710 | 0.7460594 |
| sc-3_D | Panthera spelaea | Felidae | Panthera (Panthera leo/tigris/unc | 780 | 0.74891401 |
| sc-4_O | Panthera pardus | Felidae | Panthera (Panthera leo/tigris/unc | 676 | 0.76796725 |
| sc-4_D | Panthera pardus | Felidae | Panthera (Panthera leo/tigris/unc | 927 | 0.76051209 |
| sc-5_O | Ursus spelaeus | Ursus | Ursus americanus | 1060 | 0.64648289 |
| sc-5_D | Ursus spelaeus | Ursus | Ursus arctos | 1265 | 0.68917733 |
| sc-6_O | Ursus thibetanus | Ursus | Ursus arctos | 1122 | 0.59233089 |
| sc-6_D | Ursus thibetanus | Ursus | Ursus (Ursus arctos, Ursus ameri | 1289 | 0.6098373 |
| sc-7_O | Ursus thibetanus | Ursus | Ursus arctos | 929 | 0.68166827 |
| sc-7_D | Ursus thibetanus | Ursus | Ursus arctos | 1177 | 0.6212787 |
| sc-8_O | Mammuthus sp | Proboscidea | Elephas maximus indicus | 717 | 0.72438125 |
| sc-8_D | Mammuthus sp | Proboscidea | Elephas maximus indicus | 989 | 0.60803926 |
| sc-9_O | unkown | Bovinae | Bos taurus | 699 | 0.73588593 |
| sc-9_D | unkown | Bovinae | Bison bison bison | 688 | 0.82219001 |
| sc-10_O | unkown | Ursus | Ursus arctos | 849 | 0.68490165 |
| sc-10_D | unkown | Ursus | Ursus arctos | 965 | 0.72154723 |

**Supplementary data 3: Classicol output overview table.**
Here a partial capture of the ‘benchmark_outputs.xlsx’ file is shown. This document contains the public data original file names as found in the respective papers alongside a morphological estimate of the species as provided in their supplementary data. Additionally, the proteomics outcome of each of the papers can be found next to the ClassiCOL output species. The total number of unique peptides in the sample and the ClassiCOL score can be found in this document too. In this same document in the tab ‘BoneDust output’ both the morphological output and the Classicol output of the in-house processed samples can be found, alongside any other individual sample information.

**Supplementary data 4: Detailed step-by-step protein extraction protocol,**
Product numbers are depicted in the main text.

**a. <u>Demineralization</u>**

1. Start:
   a) If bone powder: Take 5mg bone powder
   b) If bone powder on swab:
      a. Incubate while shaking 750rpm for 30min
      b. Centrifuge 15.000G 10min
      c. Discard supernatant
   c) Proteins in solution
      a. Skip to step 4
2. 600µl 5% HCl at room temperature 750rpm 24h
3. Centrifuge 10.000G 5min -> keep pellet fraction separately
4. Add 1/3 v:v TCA to the supernatant fraction (about 200µl)
5. Incubate 1h in ice
6. Centrifuge 15min 15.000G
7. Discard supernatant
8. Wash both fractions with 500µl ice cold acetone
9. Centrifuge 10min 15.000G

**b. <u>Solubilize/Reduce/Alkylate</u>**

1. Dry the pellets
2. Add 11.5 µl MilliQ water and **11.5µl lysis buffer** (SDS)
3. **Vortex**, **sonicate** for 5 min and **vortex** again
4. As needed, clarify sample of debris by centrifugation
   Centrifuge 5min @16000 g and replace the supernatant to another Eppendorf Tube
5. Add **1µl reductant** (500mM DTT)
6. Incubate **30min @37°C** in the dark
7. Add **1µl MMTS** (500mM MMTS in isopropanol)
8. **Incubate 10min @RT in the dark**
9. Add **2.5µl acidifier** and **vortex (protein crash)**
   Phosphoric acid diluted to 27.5% with water.
   E.g. 324 µL of 85% phosphoric acid diluted into 676 µL of water.
10. Check the **pH**: must be <1

**c. <u>Trap protein</u>**

11. Make binding/washing buffer
    Final concentration is 100mM TEAB in 90% Methanol. Dilute 1M TEAB with MeOH: e.g. to 1 mL 1 M TEAB, add MeOH until the final volume is 10 mL.
12. Add **165µl binding/washing buffer** to the sample
13. Bring columns on a **2ml Eppendorf** and label
14. Add sample on the column
15. Centrifuge **30sec @4000g**
    Clogged columns may be centrifuged as high as 15,000 g.
16. Reload the flow-through
17. Add **150µl washing buffer** and centrifuge **30sec @4000g.**

For best results, rotate the S-Trap micro units (like a screw or knob) 180 degrees between the centrifugations of binding and wash steps. This is especially important when using a fixed-angle rotor because the spin column does not experience homogenous flow. A mark on the outside edge during centrifugation makes it easy to track rotations.

18. Add **150µl washing buffer** and centrifuge **30sec @4000g**
19. Add **150µl washing buffer** and centrifuge **30sec @4000g**
20. Centrifuge **1min @4000g** to dry the filter
21. Place the filters on **new 1.5ml Eppendorf** tube

**d. Incubate and digest protein**

22. Make Tryp/Lys solution in 50mM TEAB: **1µg/40µl** (=20µg/800µl)
    Concentration can be adjusted to the sample, but must always be in a V of 40µl, for the blue cartridges.
23. Add **40µl** directly on the column. Remove air bubbles.
    - **Do not close the columns airtight!**
    - Visually confirm no air bubbles are present atop the trap
24. Incubate **overnight @37°C**

**e. Elute peptides**

25. Add 30µl 50mM TEAB, incubate 1min, centrifuge 1min @4000g
26. Add 30µl 0.1% FA, incubate 1min, centrifuge 1min @4000g
27. Add 30µl 50% ACN, incubate 1min, centrifuge 1min @4000g
28. Vacuum dry the sample

**Buffer and Solution Preparations:**
**2x SDS lysis buffer:**
(10% SDS + 100mM TEAB)
15ml solution made:
1.5g SDS + 1.5ml TEAB and brought to volume with Milli-Q water
**MMTS (Alkylator)**
500mM (0.5M) MMTS in isopropanol (500ul solution = 0.0005dm$^3$)
RMM MMTS = 126.20 gmol$^{-1}$ Denisity 1.337g/ml
Mols = 0.5M x 0.0005dm$^3$ = 0.00025 mols
Mass (g) = 126.20 gmol$^{-1}$ x 0.00025 mols = 0.03155g (31.55mg)
Volume of MMTS needed = 31.55mg/1.337mg/ul = **23.59ul** + 476.41 isopropanol.
**Acidifier:**
85% Phosphoric acid diluted to 27.5% with Milli-Q water. 324ul of 85% phosphoric acid diluted with 676ul of milli-Q water.
**Binding/wash buffer:**
Final concentration if 100mM TEAB in 90% methanol.
Dilute 5ml 1M TEAB with 45ml MeOH (final solution volume = 50ml)
